# Passive acoustic monitoring of Ensiferan calling diversity in a sub-tropical forest of Northeast India

**DOI:** 10.64898/2026.06.14.732046

**Authors:** Aarini Ghosh, Jishnu Borgohain, Ribankhiahbha M. War, Bittu Kaveri Rajaraman

**Affiliations:** Trivedi school of Bioscience, Ashoka University, Sonipat, 131029, India; Department of Psychology, Ashoka University, Sonipat, 131029, India; Department of Environmental Studies, Ashoka University, Sonipat, 131029, India; Foundation for Ecological Security, Shillong, 793003, India

**Keywords:** Acoustic community, Acoustic space use, tropical soundscape, Meghalaya, seasonal acoustic assemblage

## Abstract

Ensiferans are nocturnal insects (Order Orthoptera) that produce mating advertisement calls using stridulatory organs on modified forewings. These calls, typically made by males, are species-specific and serve as indicators of forest health. In biodiverse ecosystems like the subtropical forests, caller density is high, and ecological constraints such as intra- and interspecific acoustic competition, masking interference, and predation pressure can influence calling behavior. These pressures lead to variation in call structures and differences in spatiotemporal acoustic space use, leading to variations in community call type composition across the seasons. Passive Acoustic Monitoring (PAM), a non-invasive and cost-effective technique, is widely used for long-term monitoring in vertebrate taxa, but is less commonly applied to terrestrial invertebrates. In this study, we employed PAM to quantify acoustic diversity, acoustic space use, separation of different call types, seasonal calling patterns, and seasonal variation in call type composition among nocturnal Ensiferan callers. Year-round recordings were conducted using AudioMoth devices in the Khasi Hills of Meghalaya, part of the Indo-Burma Biodiversity Hotspot, at a 48 kHz sampling rate. Acoustic samples were processed using Raven Pro software. We identified 33 distinct call types, mutually distinctly differing in spectral, temporal, or both parameters. Principal Component Analysis revealed fine-scale separation of call types. While there were seasonal shifts in call types, with the dry season having the least number of callers, overall call type composition remained stable across pre-monsoon and monsoon seasons. Acoustic Space Use (ASU) analysis indicated greater use of lower frequency bands consistent with ground cricket presence, as well as seasonal variation in spectral occupancy. This foundational study is the first of its kind in Northeast India and demonstrates the potential of PAM in studying invertebrate soundscapes.

## Introduction

Insects such as Ensifera (bushcrickets and crickets) are key contributors to tropical soundscapes, especially in tropical and sub-tropical evergreen forests, where they play significant ecological roles. Their varied call structures and frequencies create acoustic diversity, which is crucial for maintaining effective communication in species-rich environments. Ensiferan call attributes such as call frequency and temporal patterns are influenced by ecological functions such as mating and anti-predator defense (Greenfield, 2002). When multiple species share the same acoustic space, interspecific acoustic interference can disrupt communication, masking and thereby reducing the effectiveness of signals. This ecological pressure can lead to species evolving distinct spectral or temporal call characteristics that are recognizable by conspecifics, with this acoustic divergence minimizing acoustic overlap and interference (Jain & Balakrishnan., 2012). For example, differences in calling frequency, timing, or duration between sympatric species reduce masking and improve mating success (Farina & Pieretti, 2014).. This divergence can also facilitate reproductive isolation, laying the groundwork for speciation (Gerhardt & Huber, 2002; (Patten *et al*. 2004). These adaptive changes often emerge in dense soundscapes like subtropical forests, where high biodiversity leads to acoustic crowding.

The concept of Acoustic Space Use (ASU) provides a framework to measure the range and overlap of frequencies used by calling species, offering insights into how they partition the soundscape (Aide *et al.,* 2017). Seasonal analysis of ASU reveals how frequency bands shift or become more crowded during periods of heightened calling activity, reflecting the influence of habitat features and environmental changes on acoustic behavior. By examining ASU across seasons and habitats, it becomes possible to identify patterns of seasonal acoustic partitioning, revealing how environmental pressures drive acoustic variation. Such analysis examines the acoustic niche hypothesis, which posits that different species partition the soundscape to avoid interference from both conspecific and heterospecific calls (Krause, 1993). These tools enable assessments of species diversity, acoustic separation, and community structure.

Passive acoustic monitoring (PAM) is a powerful, non-invasive method for studying acoustic diversity and call divergence, particularly in challenging environments like dense forests (Sugai et al., 2018). Autonomous recording devices used in PAM enable continuous data collection across temporal and spatial scales, capturing fine-scale variations in calling behavior and acoustic diversity. PAM can be a better approach to monitoring Ensiferan species, allowing researchers to document temporal and spatial patterns of calling activity across seasons and habitats (Symes *et al*. 2022). By providing a season-spanning dataset, PAM offers a comprehensive view of how environmental factors and habitat differences influence calling behavior. Despite its advantages, PAM has limitations. It primarily captures data from acoustically active species, potentially underrepresenting silent or low-frequency species. Overlapping calls and the lack of individual vocalization differentiation can pose challenges for species identification and abundance estimation. Large PAM datasets also require sophisticated storage and analysis tools, and substantial computational resources. Additionally, non-target sounds such as wind, rain, or human activity can interfere with data quality, complicating the extraction of species-specific calls. While automated species classification algorithms are improving, they often require manual verification to ensure accuracy.

Acoustic monitoring in ecological research offers numerous advantages and presents certain challenges. It enables the simultaneous study of multiple taxonomic groups using versatile recording equipment capable of capturing sounds across various frequency ranges. Temporal data collection allows the investigation of nocturnal, diurnal, monthly, and seasonal variations, even in areas that may otherwise be inaccessible. Spatial sampling can be scaled up to test ecological hypotheses across large gradients, while functional ecological links, such as interactions between sound-producing organisms and their environments, can also be explored. However, challenges include variations in species vocalizations, non-uniform signal propagation due to habitat and weather conditions, limitations in power and storage for remote setups, and interference from anthropogenic and environmental non-target sounds. These factors underscore the potential of acoustic monitoring while highlighting its logistical and analytical complexities (Fig. 4.1). (Schmidt *et al*. 2013, Ross *et al*. 2023).

In this study, we utilize PAM to investigate the processes driving call divergence in Ensiferan species. By compiling a season-spanning dataset from sub-tropical forests in Northeast India, we examine how Ensiferan call types vary across temporal and spatial scales. Long-term PAM monitoring provides the opportunity to detect gradual changes in acoustic parameters among sympatric species, potentially revealing early stages of divergence driven by evolutionary pressures within shared habitats. By integrating acoustic diversity metrics with seasonally structured datasets, this research advances the understanding of how Ensiferan species utilize sound to minimize interference and maintain distinct communication channels.

### Study sites

This study was conducted near the Nongkhyllem Forest Reserve, in the village of Umling (25°57’56.6“N, 91°51’37.9“E), located in the Ri-Bhoi District of Meghalaya (Fig. 2B.). The reserve is a highly diverse subtropical forest within the Indo-Burma biodiversity hotspot of Northeast India (Fig 1). Field sound recordings were carried out in Umling, where two transect sites were selected (Fig. 2A). Both sites are situated near human settlements, with vegetation consisting of a mix of forests and plantations. The vegetation structure includes grass understory, middle story, and canopy layers. For transects site 1 was in a small hilly region where altitude changes at every point and site 2 had had almost a constant altitude of 300 m (a.m.s.l.). The elevation across site 1 ranges from 250 to 500 meters above the mean sea level (a.m.s.l.). The region experiences an average annual temperature between 15°C and 28°C.

**Figure 1.**
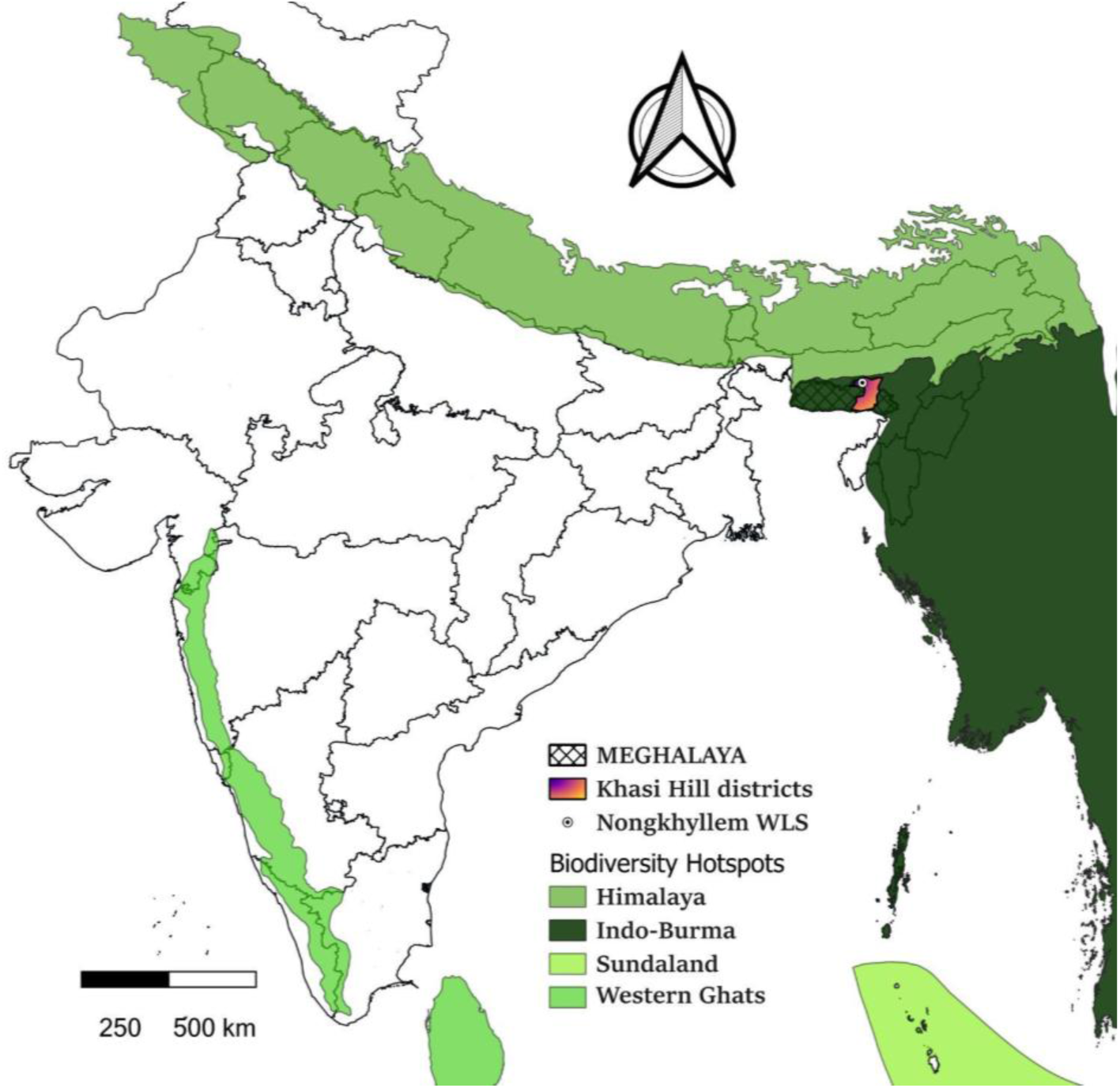
Map showing the study site Nonkhyllem WLS in India with the state boundary of Meghalaya and distribution of Khasi Hill districts including Ri-Bhoi. Besides that distribution of four different Biodiversity hotspots of India shown in green colour gradients.

**Figure 2.**
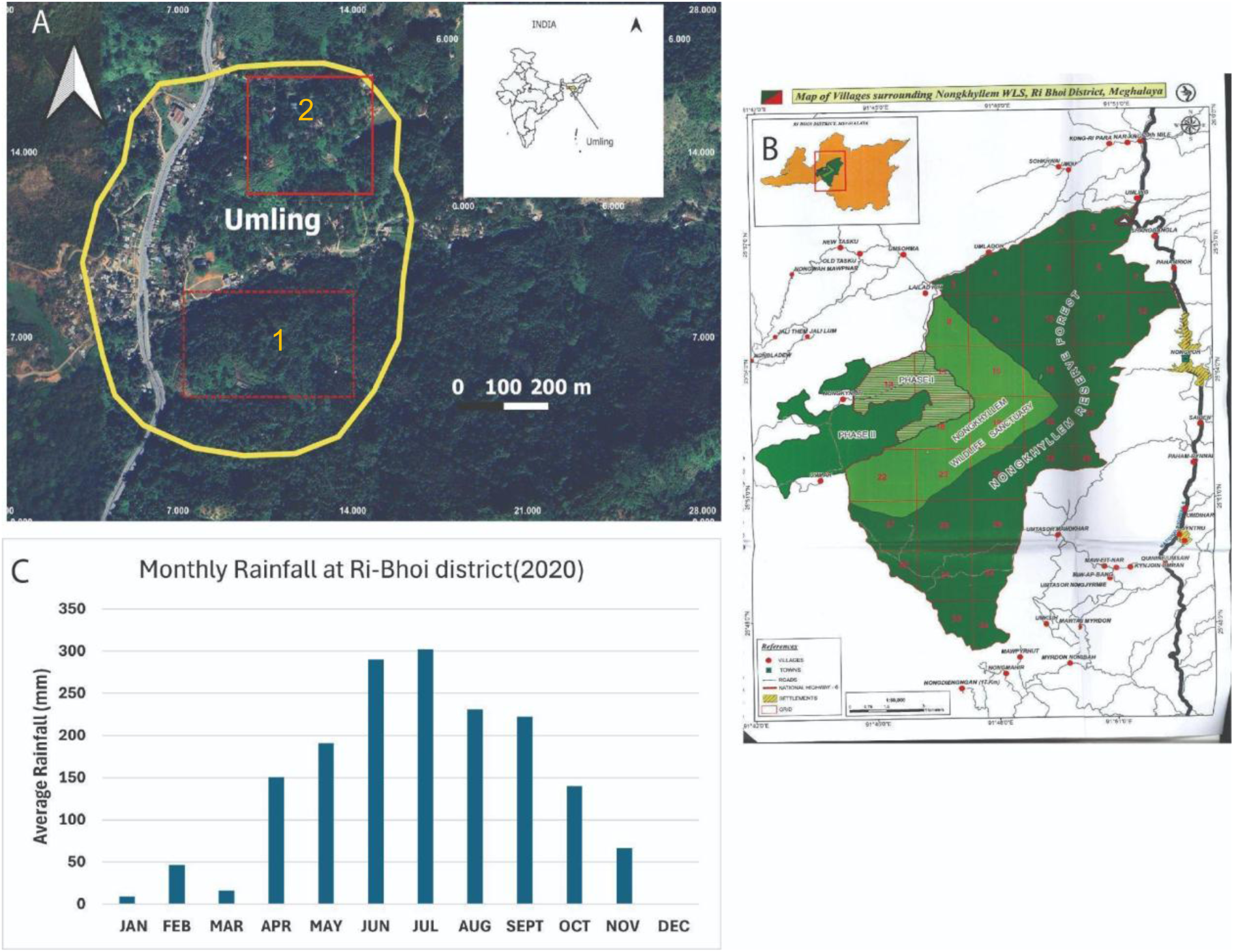
Maps of study sites and average rainfall data (A) Google Street view of Study Sites Using Google Earth with the area of both transect sites. The dotted red line denotes locations for transect site 1 and solid red lines for transect site 2 (B) The boundary of Nongkhyllem forest reserve and the location of our study site Umling village (Map from Meghalaya Forest department) (C) Average rainfall at Ri-Bhoi district every month. (data from: www.megagriculture.gov.in)

Meghalaya receives high rainfall, with significant seasonal variations. In Ri-Bhoi district during the dry months, rainfall averages as low as 10 mm, while in the monsoon season, it can reach up to 350 mm, based on data from the Meghalaya State Agricultural Department (Fig. 2C). The seasons were categorized into four groups based on rainfall patterns:

1. Pre-monsoon (February to April): Medium rainfall, averaging 30–150 mm.
2. Monsoon (May to July): Highest rainfall, averaging 200–300 mm.
3. Post-monsoon (August to October): Moderate rainfall, averaging 140–230 mm.
4. Dry season (November to January): Lowest rainfall, averaging 0–60 mm.

### Material and Methods

## Acoustic sampling and recording

Acoustic sampling was conducted using passive acoustic recorders along two 500-meter transect sites (Fig. 1A). Each transect was surveyed twice in a month, with the transect line divided into 10 equidistant points at intervals of 50 meters each. Audio recordings were captured using AudioMoth 2.0 devices for 12 minutes per point at a sampling rate of 48 kHz. Recorders were positioned at a height of 6–7 feet and secured horizontally to a tree or twig. Figure. 3 illustrates the detection capabilities and inherent limitations of the AudioMoth recorder.

**Figure 3.**
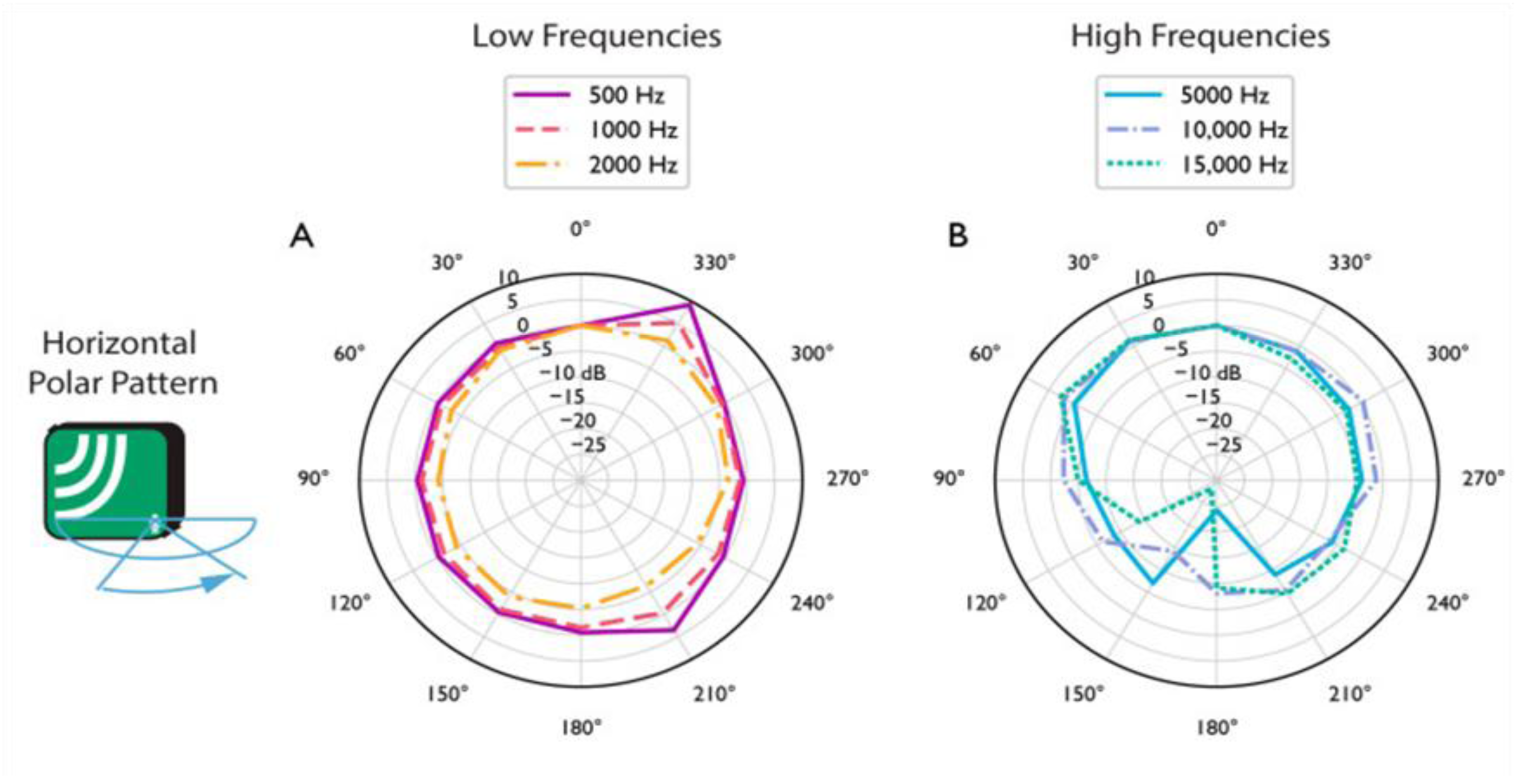
Detectability of different frequencies by the AudioMoth device (Picture ref: Lapp *et al.,* 2023).

Sampling was conducted over nine months, excluding the monsoon season, as heavy nocturnal rainfall during this period masked biophony. Recordings were performed twice per month at each transect site, typically on no-moon nights between 8:00 PM and 12:00 AM, corresponding to the peak activity period of Ensiferans. All recordings were done over the span of a year from August 2022 to April 2023. While the primary focus of the recordings was to capture Ensiferan calls, the passive acoustic method precluded taxonomic identification of the callers.

To address this limitation, psychoacoustic sampling was employed to validate the passive acoustic recordings. This approach enabled the identification of four Ensiferan species, two at the species level and two at the genus level. However, the identification of additional callers was hindered by the lack of a comprehensive, open-access database for Ensiferan calls specific to the region. Accumulation curves for call types in each site were made to show the completeness of all call types in this particular location.

### Acoustic analysis

A total of 360 recordings, each 12 minutes in duration, were collected over nine months (40 recordings per month). All recordings were analyzed using Raven Pro software. Call types were initially differentiated by visualizing their distinct frequency patterns in spectrograms through the FFT (fast fourier transform) function (Fig. 4). Each distinct call type was categorized and annotated using psychoacoustic methods in Raven Pro 1.6 and assigned an arbitrary caller identification number (ID) to group similar calls that seemed to fall within a single call type.

The temporal and spectral characteristics of each call type were analyzed in Raven Pro, with annotations cleaned and refined in Audacity 3.0. Temporal parameters measured included echeme period, syllable period, and syllable repetition rate, while spectral parameters included peak frequency and bandwidth. A standardized protocol, as outlined in the methods of the first chapter, was employed for these analyses. MATLAB (R2022a) was used to generate all oscillograms and spectrograms for each call type, providing a detailed visualization of their acoustic characteristics. All the temporal and spectral parameters of all calls which were recorded at different temperatures regressed against the optimal temperature of 24°C. The normality of the data was assessed using the Shapiro Wilk’s test. As the data were normally distributed, the mean and standard deviation of each acoustic parameter were calculated for further analysis.

### Acoustic community assemblage

To observe the nocturnal acoustic community structure, we plotted the call syllable repetition rate as a measure for temporal variation against call peak frequency for spectral variation (referencing Diwakar & Balakrishnan, 2007), year-round and in different seasons. Scatter plots were made in GraphPad Prism 2 software.

### Acoustic separation of each call type using principal component analysis (PCA)

We used a hierarchical approach to examine acoustic separation between call types based on acoustic characteristics. We used spectral and the short time-scale temporal characters (syllable and echeme) but not any large-scale temporal separation information (seasonal or diel activity) for this. All call types were first visualized by plotting their peak frequency against their bandwidth since in the spectrogram already these call types show relatively clear separation. Then sub-categorized the callers based on their spectral characteristics as either low-frequency, high-frequency narrow-band, or high-frequency broad-band callers. Triller call types were also separated from chirpers and complex calls based on their unique temporal patterns. PCAs were then performed in RStudio 2023 (version 06.01) within each category using quantified temporal characters to visualize the separation of call types.

### Acoustic space use and nocturnal bioacoustic diversity calculation

Acoustic Space Use (ASU) was calculated for the identified call types in this particular region of forest for the specific time of 8 PM-12 AM in each transect site throughout the seasons. The ASU was calculated relative to 24 bins of 1 kHz bin size, with the Nyquist frequency constrained by the sampling rate of 48kHz. The proportion of frequency-band use in each bin was calculated as the number of callers using that bin, normalized by 11 - the maximum value reached in any frequency bin. Each caller’s contribution was considered to be only in the bin containing the peak frequency, for all narrow-band callers. The contribution of all broadband callers was considered to be the full bandwidth: that is the range between the maximum and minimum frequency with power within 20db of the power at the peak frequency. A heat map was then generated in RStudio using the proportion of frequencies in each bin to visualize the ASU.

To calculate the proportion of acoustic space used for each season, we first sum the ASU values across all frequency bins used in that season. This total ASU is then divided by the total number of available frequency bins, and the result is multiplied by 100 to express the proportion as a percentage:

**The proportion of ASU** = (Seasonal ASU / Total number of bins) × 100

This calculation provides a relative measure of acoustic space occupancy, enabling seasonal comparisons of how much of the acoustic frequency range is actively used by all call types, a method commonly applied in eco-acoustic studies (e.g., Aide *et al*. 2017., Staniweicz *et al*. 2023).

### Call occurrence analysis and seasonal variation

We analyzed the occurrence of different call types using two approaches. First, Wecalculated the binary presence/absence of each call type (absence was denoted as 0 and presence as 1) for the passive recording made at each grid point within a transect. However, it remains uncertain whether the same individual’s call was recorded multiple times across different points within a transect, or whether each data point represents a different caller, as no physical evidence of the caller’s movement was available. This occupancy analysis focused solely on the presence or absence of each call type. From this, we calculated the occurrence rate as the total number of detections of each call type at a site divided by the total number of grid points at that site. Subsequently, we averaged the data from two repeated transects across all months within a season to derive the seasonal occupancy rate. We generated heat maps based on these occupancy rates to visualize the distribution and seasonal occurrence of each call type across different transects.

Secondly, we examined the variation in the occurrence rates of each call type across seasons, as well as the overall seasonal variation in the occurrence of multiple call types. This approach provided insights into seasonal patterns and variations in call type occurrence.

The occurrence rate was calculated with the following formula:

**Occurrence Rate** = (Number of segments where the caller is detected) / (Total number of segments in the season)

### Multivariate analysis of call type compositions in each season

The variation in call type composition in each season at both sites was visualized using the nonparametric multivariate tool NMDS (nonlinear multidimensional scaling) on call type occurrence data in RStudio.

### Statistical analysis

To examine the variation across seasons, the Shapiro test was done to check normality. When the data were normal, an ANOVA and Tukey’s test were done; else Kruskal-Wallis’s test and Dunn’s test were done, and the data were represented by medians and interquartile ranges (IQR).

A generalized linear model (GLM) was used to study the difference in the variation in occurrence rate in each season. The model was fitted by restricted maximum likelihood (REML) and used Satterthwaite’s approximation to calculate degrees of freedom.

An ANOVA was done with the regressed axis of NMDS 1 and NMDS 2 values to understand the statistical difference in call type composition across seasons for both transects. All statistical tests were performed with RStudio-2023 06.01 software.

## Results

### 1. Accumulation curve of call types

Each pair of consecutive days was sampled from each month, and the curve represents a year’s worth of data, where the rainy season data do not add species identity information to the sampling due to the sound of rain masking all other acoustic information. (Fig. 5)

**Figure 4.**
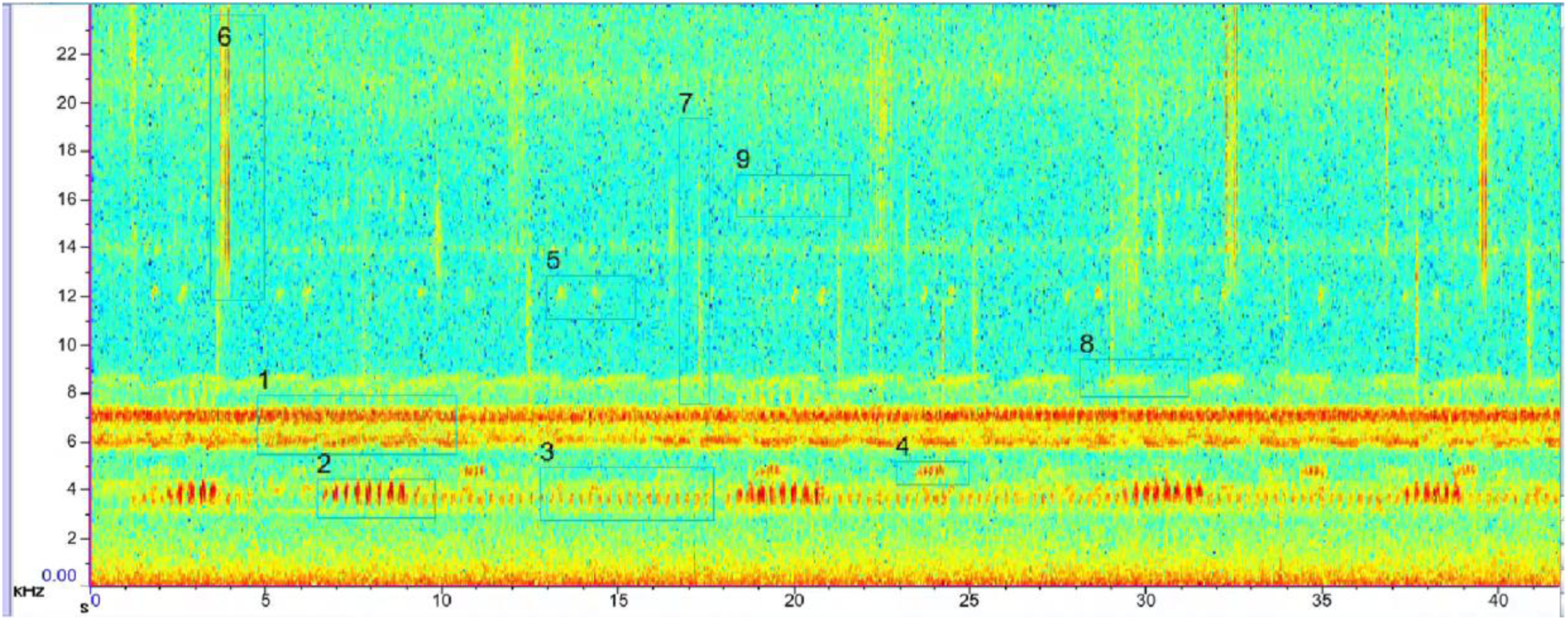
A passive acoustic recording spectrogram from the field. Each numeric number on the spectrogram next to a box shows all different call types that can be extracted from a single passive recording.

**Figure 5.**
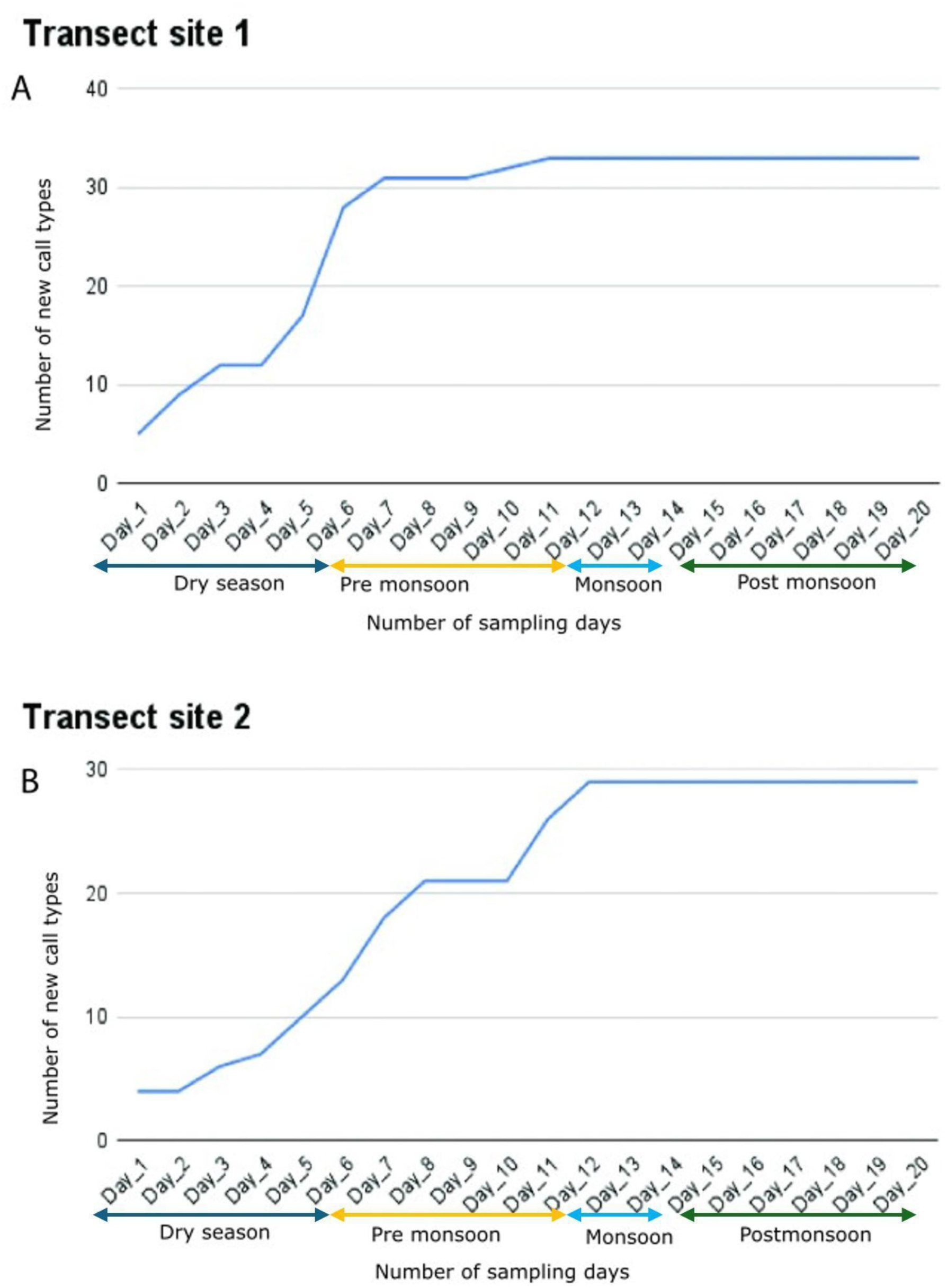
Call type accumulation curves across days of passive acoustic sampling from two different transect sites. Each season consists of 3 months, with 2 collection dates per month.

This shows that the species accumulation curve flattens out after the 11th day for data collected from transect 1 and on the 12th day for transect 2 of the survey. This curve was made based on passive recording data from each day. The first 6 days are from the Dry season, the 7th to 12th days are from the Pre Monsoon season, the 13th and 14th from the Monsoon (where the sound of the rain masked all other acoustic information), and the 15th to 20th are from the Post Monsoon season.

#### 2. Detailed spectral and temporal analysis of all callers

After thoroughly analyzing the FFT of all recordings manually over the course of a year, a total of 33 distinct call types were identified and annotated. These call types are not only psychoacoustically distinct but also exhibit clear differences in their temporal and spectral parameters. Each call type was assigned an arbitrary identification number (ID) for reference. Audio recordings are available here https://doi.org/10.5281/zenodo.14636913. All spectrograms and oscillograms are attached in the supplementary material (Figure S1- S4).

#### ID-1

This call consists of a continuous trill with indistinct inter-syllable duration. The syllable period is 0.004 +/- 0.001 s (mean +/- SD, n = 50). This broadband call is present across the span of the recorded spectrum with peak frequency and bandwidth of 6.46 +/- 0.5 kHz and 10.12 +/- 1 kHz (median +/- IQR, n = 5) respectively.

#### ID-2

The caller has a series of distinct echemes of 0.17 +/- 0.1 s (mean +/- SD, n = 50). Each echeme consists of syllables with a period of 0.02 +/- 0.001 s (mean +/- SD, n = 50). This is a narrowband call with peak frequency and bandwidth of 3.95 +/- 0.05 kHz and 1.45 +/- 0.28 kHz (median +/- IQR, n = 5) respectively.

#### ID-3

This is a chirper with distinct echemes with a period of 0.81 +/- 0.17 s (mean +/- SD, n = 10). Echemes had a syllable period of 0.04 +/- 0.03 s (mean +/- SD, n = 50). This is a narrowband call with a peak frequency of 4.7 +/- 1.2 kHz (mean +/- SD, n = 5), and a bandwidth of 0.5+/-0.2 kHz (median +/- IQR, n = 5).

#### ID-4

This is a very low-duty cycle call with a single echeme with a long pause with an echeme period of 2.8 +/- 2 s (mean +/- SD, n = 50). The echeme has only two syllables with a syllable period of 0.11 +/- 0.06 s (mean +/- SD, n = 50). The peak frequency is 11.6 +/-0.9 kHz (median +/- IQR, n=5), and the bandwidth is 9+/-0.11 kHz (median +/- IQR, n = 5).

#### ID-5

This chirper call type has a distinct echeme with an echeme period of 0.2 +/- 0.2 s (mean +/- SD, n = 50) and a syllable period of 0.05 +/- 0.01 s (mean +/- SD, n=50). The peak frequency is 12.3 +/- 0.3 kHz (median +/- IQR, n = 5), and the bandwidth is 0.9 +/- 0.04 kHz (median +/- IQR, n = 5).

#### ID-6

This high-frequency broadband call has a very distinct high-duty cycle echeme. Echeme period is 0.06 +/- 0.05 s (mean +/- SD, n=50) and syllable period is 0.001 s. The peak frequency and bandwidth are 21.1+/-1.5 kHz (median +/- IQR, n = 5) and 41.6 +/- 0.9 kHz (median +/- IQR, n = 5) respectively. This caller was taxonomically identified as *Ducetia sp*.

#### ID-7

This caller was taxonomically identified in the field as *Mecopoda “Complex”* call type. This is a broadband loud caller with a verse of three different segments. The first two segments have distinct echemes and the end segment is a continuous trill. Detailed call features of this call are already described in Chapter 2.

#### ID-8

This broadband high-frequency caller was identified in the field and characterized morphologically and taxonomically as *Hexacentrus unicolor.* Their call has an echeme period of 0.118 +/- 0.04 s (mean +/-SD, n =5 0) and a syllable period of 0.003 s. The peak frequency and bandwidth are 13 +/- 0.5 kHz (median +/- IQR, n = 5) and 39.5 +/- 1.2 kHz (median +/- IQR, n = 5) respectively.

#### ID-9

This loud ground cricket caller has a distinct echeme with an echeme period of 0.5 +/- 0.26 s (mean +/-SD, n = 60) and a syllable period of 0.003 s. This low-frequency caller was taxonomically identified up to the genus level as *Gymnogryllus sp.* The peak frequency and bandwidth are 3.85+/- 0.05 kHz (median +/- IQR, n = 6) and 1.3 +/- 0.1 kHz (median +/- IQR, n = 6), respectively.

#### ID-10

This triller has a syllable period of 0.009 s (mean +/- SD, n = 10) with a peak frequency of 4.4 kHz and a narrow bandwidth of 1.5 kHz (median, n = 2) .

#### ID-11

This is another triller with a peak frequency of 4.3 +/- 0.28 kHz (median +/- IQR, n = 2) and bandwidth 0.8kHz. The syllable period is 0.015 +/- 0.08 s (mean +/- SD, n = 20).

#### ID-12

This chirper has an echeme period of 0.25 +/- 0.07 s (mean +/- SD, n = 50) and syllable period of 0.066+/- 0.1s (mean +/- SD, n = 50), with a peak frequency of 3.28 +/- 0.21 kHz (median +/- IQR, n = 5) and bandwidth of 1.2 +/- 0.4 kHz (median +/- IQR, n = 5).

#### ID-13

This is a narrow band caller with an echeme period of 0.113 +/- 0.04 s (mean +/- SD, n = 20) and a syllable period of 0.017 +/- 0.001 s (mean +/- SD, n = 20). The peak frequency and bandwidth are 7 kHz and 0.5 kHz (median, n = 2) respectively.

#### ID-14

This triller has a syllable period of 0.1 +/- 0.08 s (mean +/- SD, n = 50) of peak frequency 2.94 +/-1.6 kHz (median +/- IQR n = 5) and bandwidth 0.7 +/- 0.01 kHz (median +/- IQR, n = 5).

#### ID-15

This triller shows amplitude modulation in the call. The syllable period is 0.02 s with a peak frequency of 8.3 +/-0.2 kHz (median +/- IQR, n = 5), and a bandwidth of 2 +/- 0.04 kHz (median +/- IQR, n = 5).

#### ID-16

The echeme consists of two syllables. Syllable and echeme periods are 0.02 +/- 0.02 s (mean +/- SD, n = 40) and 0.13 +/- 0.12 s (mean +/- SD, n=40), respectively. The peak frequency and bandwidth are 4.5 +/- 0.3 kHz (median +/- IQR, n = 4) and 1.8 +/- 0.12 kHz (median +/- IQR, n = 4).

#### ID-17

This is a chirper with an echeme period of 3.4+/-1.17 s (mean +/- SD, n = 30) and syllable period 0.28 +/- 0.01 s (mean +/- SD, n = 30). The peak frequency and bandwidth of this narrow-band call are 3.9 +/- 0.9 kHz (median +/- IQR, n = 3) and 0.63+/-0.14 kHz (median +/- IQR, n = 3).

#### ID-18

This chirper has an echeme period of 0.4 +/- 0.04 s (mean +/- SD, n = 40) and a syllable period of 0.06 +/- 0.04 (mean +/- SD, n = 40). The frequency bandwidth and peak frequency are 0.75 +/- 0.1 kHz and 7.7 +/- 0.05 kHz (median +/- IQR n = 4) respectively.

#### ID-19

This is a low-duty cycle caller with an echeme period of 0.2 +/- 0.1 s (mean +/- SD, n = 40) and a syllable period of 0.006 +/- 0.003 s (mean +/- SD, n = 40). Peak frequency and bandwidth are 5.4 +/- 0.4 kHz and 2.3 +/- 0.5 kHz (median +/- IQR, n = 4), respectively.

#### ID-20

The call consists of an echeme period and syllable period of 0.8 +/- 0.33 s (mean +/- SD, n = 20) and 0.03 +/- 0.01 s (mean +/- SD, n = 20), respectively. The peak frequency is 5.56 +/- 0.5 kHz (median +/-IQR, n = 2) with a bandwidth of 0.6 +/- 0.1 kHz (median +/- IQR, n = 2).

#### ID21

This is a chirper with an echeme period of 0.5 +/- 0.5 s (mean +/- SD, n = 30) and syllable period of 0.05 +/- 0.01 s (mean +/- SD, n = 30). Peak frequency and bandwidth are 5.4 +/- 0.2 and 2 +/-0.5 kHz (median +/- IQR, n = 3) respectively.

#### ID-22

This is a high-duty cycle chirper with an echeme period and syllable period of 8.1 +/- 7.1 s and 0.025 +/- 0.013 s (mean +/- SD, n = 40) respectively. The peak frequency is 6.95 +/- 0.12 kHz (median +/-IQR, n = 4), and the bandwidth is 2 +/- 0.2 kHz (median +/- IQR, n = 4).

#### ID-23

This is a chirper with an echeme period of 1 +/- 0.4 s (mean +/- SD, n = 50) and syllable period of 0.05 +/- 0.01 s (mean +/- SD, n = 50). The peak frequency and bandwidth are respectively 4.6 +/- 0.4 kHz and 1.18 +/- 0.1 (median +/- IQR, n = 5).

#### ID-24

This triller call type has two segments in each verse. The first segment (ID-24_I) is of high amplitude syllables and the second (ID-24_II) is of low amplitude syllables. The first segment has a syllable period of 0.028 +/- 0.01 s (mean +/- SD, n = 50), and the second segment has a syllable period of 0.023 +/- 0.01 s (mean +/- SD, n = 50). However, there is no difference in spectral parameters between both segments. Peak frequency and bandwidth are 2.4 +/- 0.2 kHz (median +/- IQR, n = 5) and 0.4 respectively.

#### ID-25

This continuous triller has a syllable period of 0.1 +/- 0.02 s (mean +/- SD, n = 50), a peak frequency of 2.74 +/- 0.13 kHz (median +/- IQR, n = 5) and a bandwidth of 0.2 kHz.

#### ID-26

This is a low-duty cycle broadband call. The echeme period and syllable period are 0.2 +/- 0.003 s (mean +/- SD, n = 30) and 0.05 +/- 0.002 s (mean +/- SD, n = 30) respectively. The peak frequency is 4.8 +/- 0.1 kHz and the bandwidth is 6 +/- 0.3 kHz (median +/- IQR, n = 3).

#### ID-27

This is another low-duty cycle low-frequency caller with an echeme period of 0.5 +/- 0.5 s (mean +/-SD, n = 20) and syllable period of 0.5 +/- 0.55 s (mean +/- SD, n = 20). The peak frequency and bandwidth are 2.3 +/- 0.14 kHz (median +/- IQR, n = 2) and 1.6 +/-0.1 kHz (median +/- IQR, n = 2) respectively. This sounds as if it might be an anuran call.

#### ID-28

This high-frequency chirper has an echeme period of 1 +/- 0.5 s (mean +/- SD, n = 30) and syllable period of 0.05 +/- 0.01 s (mean +/- SD, n = 30). The peak frequency and bandwidth are 11.4 +/- 0.1 kHz (median +/- IQR, n = 3) and 2.4 +/- 0.1 kHz (median +/- IQR, n = 3) respectively.

#### ID-29

A high-duty cycle chirper with echeme period of 2 +/- 0.1 s (mean +/- SD, n = 20) and syllable period of 0.006 +/- 0.002 s (mean +/- SD, n = 20). The peak frequency and bandwidth are 7.8 +/- 0.7 kHz (median +/- IQR, n = 2) and 2.4 +/- 0.1 kHz (median +/- IQR, n = 2) respectively.

#### ID-30

Echeme period for this call is 0.14 +/- 0.02 s (mean +/- SD, n = 20), and the syllable period is 0.01+/- 0.001 s (mean +/- SD, n = 20). The peak frequency and bandwidth of this call are respectively 6.1 +/- 0.1 kHz (median +/- IQR, n = 3) and 7.3 +/-0.3 kHz (median +/- IQR, n = 3).

#### ID-31

This is a high-frequency triller call with indistinguishable syllables. The syllable period is 0.007 +/- 0.003 s (mean +/- SD, n = 20). The peak frequency and bandwidth are 11.1 +/- 0.14 kHz median +/-IQR, n = 2) and 2 +/-0.07 kHz (median +/- IQR, n = 2) respectively.

#### ID-32

This low-duty cycle chirper has a syllable period and echeme period of 0.04 +/- 0.002 s (mean +/- SD, n = 20) and 0.13 +/- 0.01 s (mean +/- SD, n = 20) respectively. The Peak frequency for this call is 4.8 +/- 0.1 kHz (median +/- IQR, n = 2), and bandwidth of 1.15 +/- 0.07 kHz (median +/- IQR, n = 2).

#### ID-33

This is a high-frequency narrowband chirping call with an echeme period and syllable period of 0.9 +/- 0.04 s and 0.05 +/- 0.01 s (mean +/- SD, n = 30) respectively. The peak frequency is at 17.3 +/- 0.5 kHz (median +/- IQR, n = 3), and the bandwidth is 2.1 +/- 0.1 (median +/- IQR, n = 3).

### 3. Categorization of different call types

To understand how the call types are differentiated, our first level of categorization was based on their fundamental temporal and spectral patterns. Calls exhibiting grouped syllables forming an echeme with distinct inter-echeme durations were categorized as “chirper” types. Conversely, calls lacking such grouping or distinct echeme formation were categorized as “triller” types. Among the 33 identified calls, 7 were classified as triller call types (ID-1, ID-11, ID-14, ID-15, ID-24, ID-25, and ID-31). ID-7 was identified as a complex call type, containing both trill and chirp components. The remaining call types were categorized as chirper types (Table 1).

**Table 1.** Categorized list of all call types based on broad temporal, spectral and band width.

| Call types | Broad temporal category | Broad spectral category | Band width category |
| --- | --- | --- | --- |
| ID_1 | Trill | Low frequency | Broad band |
| ID_2 | Chirp | Low frequency | Narrow band |
| ID_3 | Chirp | Low frequency | Narrow band |
| ID_4 | Chirp | High frequency | Narrow band |
| ID_5 | Chirp | High frequency | Narrow band |
| ID_6 | Chirp | High frequency | Broad band |
| ID_7 | Chirp | High frequency | Broad band |
| ID_8 | Chirp | High frequency | Broad band |
| ID_9 | Chirp | Low frequency | Narrow band |
| ID_10 | Chirp | Low frequency | Narrow band |
| **ID_11** | Trill | Low frequency | Narrow band |
| **ID_12** | Chirp | Low frequency | Narrow band |
| **ID_13** | Chirp | High frequency | Narrow band |
| **ID_14** | Trill | Low frequency | Narrow band |
| **ID_15** | Trill | Low frequency | Narrow band |
| **ID_16** | Chirp | Low frequency | Narrow band |
| **ID_17** | Chirp | Low frequency | Narrow band |
| **ID_18** | Chirp | High frequency | Narrow band |
| **ID_19** | Chirp | Low frequency | Narrow band |
| **ID_20** | Chirp | Low frequency | Narrow band |
| **ID_21** | Chirp | Low frequency | Narrow band |
| **ID_22** | Chirp | High frequency | Narrow band |
| **ID_23** | Chirp | Low frequency | Narrow band |
| **ID_24** | Trill | Low frequency | Narrow band |
| **ID_25** | Trill | Low frequency | Narrow band |
| **ID_26** | Chirp | Low frequency | Narrow band |
| **ID_27** | Chirp | Low frequency | Narrow band |
| **ID_28** | Chirp | High frequency | Narrow band |
| **ID_29** | Chirp | High frequency | Narrow band |
| **ID_30** | Chirp | Low frequency | Narrow band |
| **ID_31`** | Trill | Low frequency | Narrow band |
| **ID_32** | Chirp | Low frequency | Narrow band |
| **ID_33** | Chirp | High frequency | Narrow band |

**Table 2:**
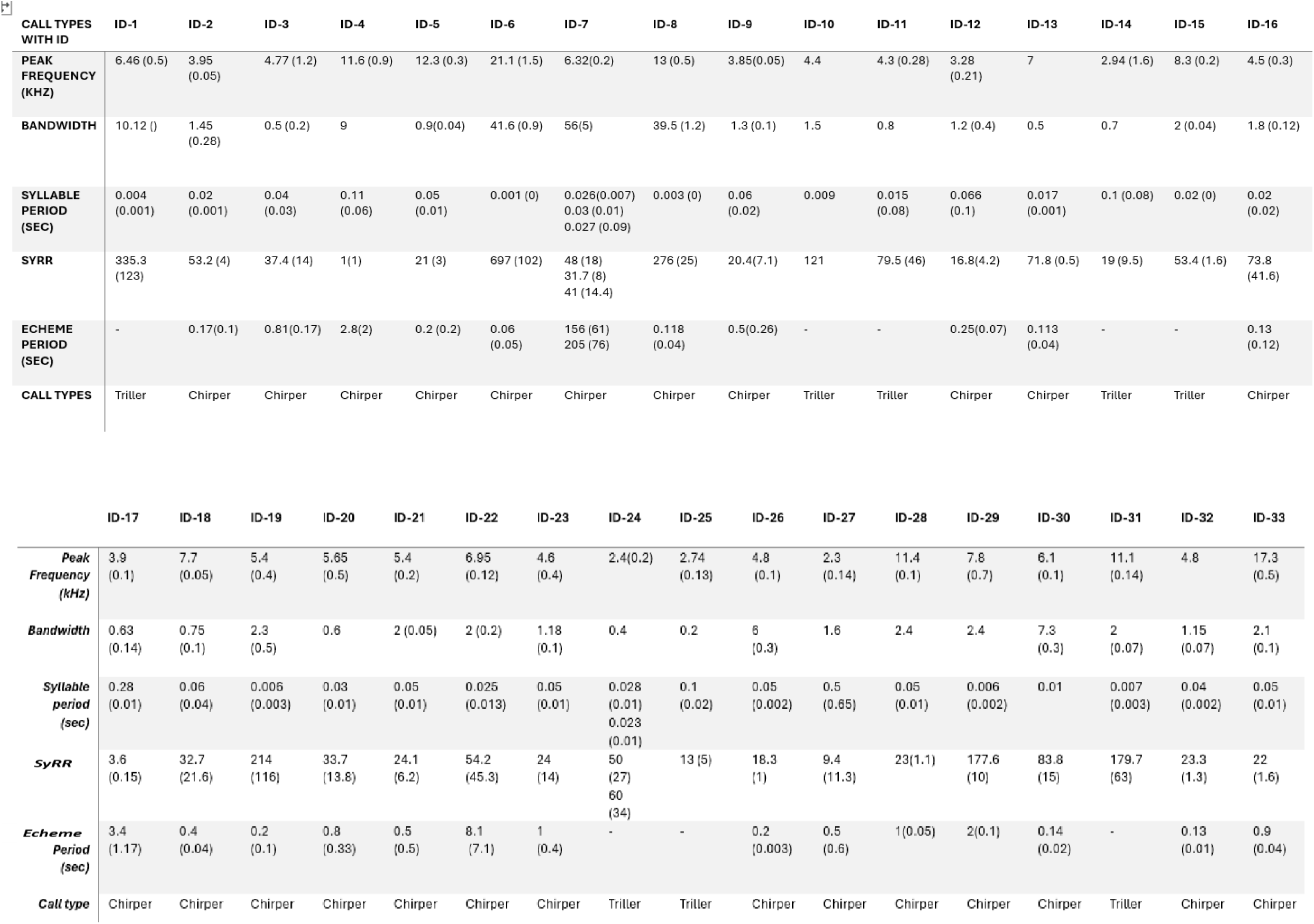
Detailed temporal and spectral characteristics of all call types. The number in the brackets denotes the standard deviation.

Each caller also exhibited unique spectral characteristics. Based on average peak frequency and bandwidth, all callers were grouped into three main categories: low-frequency callers, high-frequency narrow-bandwidth callers, and high-frequency broadband callers (Table 1). A scatter plot illustrates a clear separation between high-frequency and low-bandwidth calls, with the majority of callers falling into the low-frequency narrow-band category (Fig. 6).

**Figure 6.**
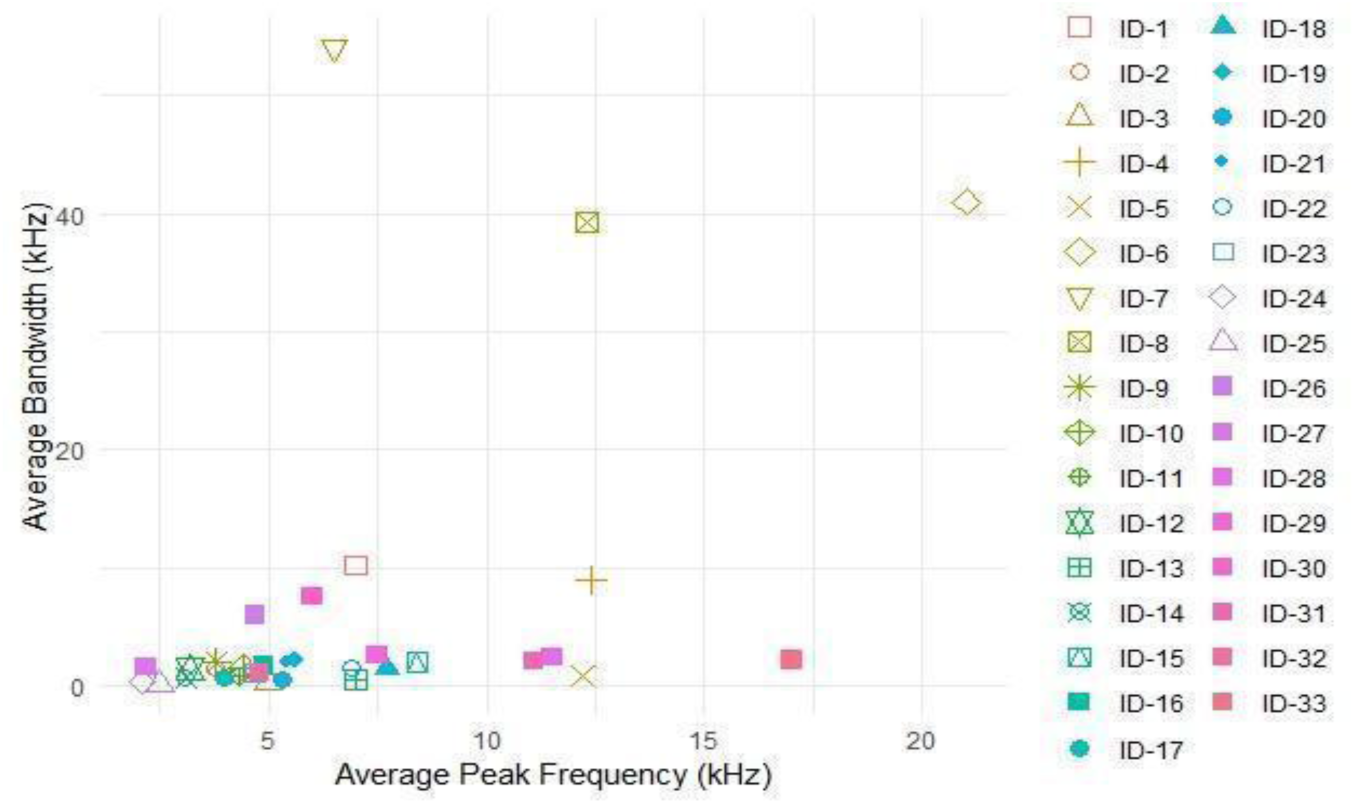
Scatter plot showing different call types based on average peak frequency and their average bandwidth. Different shaped markers with color combinations denote different call types.

### 4. Acoustic community assemblage of callers in different seasons across both sites

The nocturnal acoustic community was composed of 33 different call types, excluding all nocturnal bat calls. These recordings may not be limited to Ensiferans and might include some Anurans.

Even though there is a diverse number of call types, all callers are not present throughout all the seasons. The total number of call types changes from season to season as well as between the two transects (Fig.16).

#### 4.a. Acoustic community assemblage of callers throughout the season

The acoustic community assemblage of all the 33 different call types recorded throughout the year across all three seasons and transects (without monsoon season) covers a wide range of syllable repetition rates and frequencies (Fig. 7). A log plot was necessary to maximize separation, and the majority of the call types were found to have frequency ranges from 4-8kHz. ID24 and ID 27 have very low frequency, and they separate from the rest of the call types. ID 4 is well separated from the rest of the call types with very low SyRR. Some higher peak frequency callers like ID6, ID8 and ID3 have correlated high values of both frequency and SYRR and therefore are well separated from each other and the other call types based on their acoustic characters. The most closely overlapping call types are low frequency, low syllable repetition rate callers.

**Figure 7.**
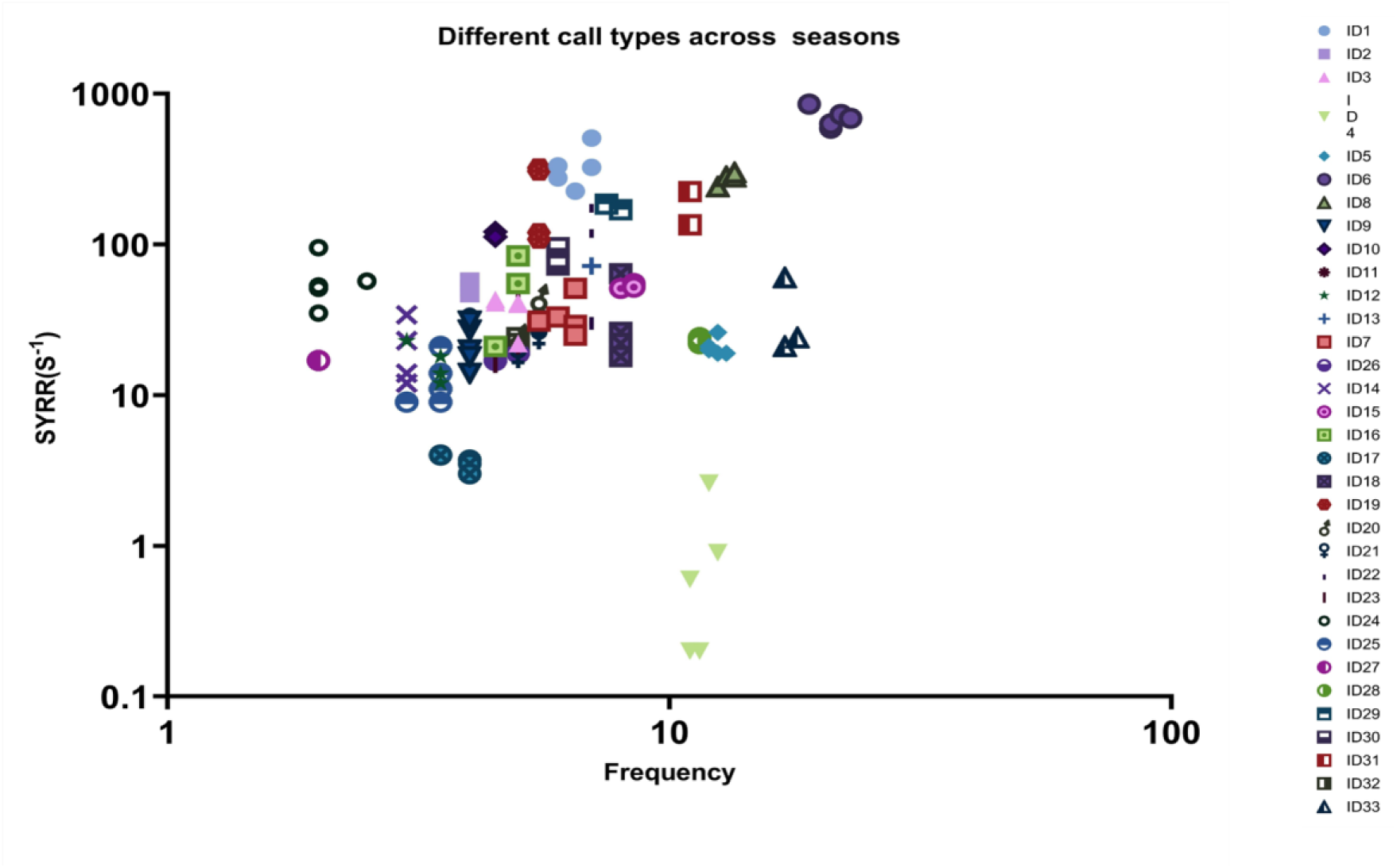
Distribution of different call types relative to syllable repetition rate (SyRR) and peak frequency (kHz) in log scale. Most of the callers are in the low-frequency and high SyRR range.

#### 4.b. Acoustic community assemblage of the callers in different seasons

Three separate scatter plots were made to understand the variation in acoustic community assemblage between seasons (Fig. 8).

**Figure 8.**
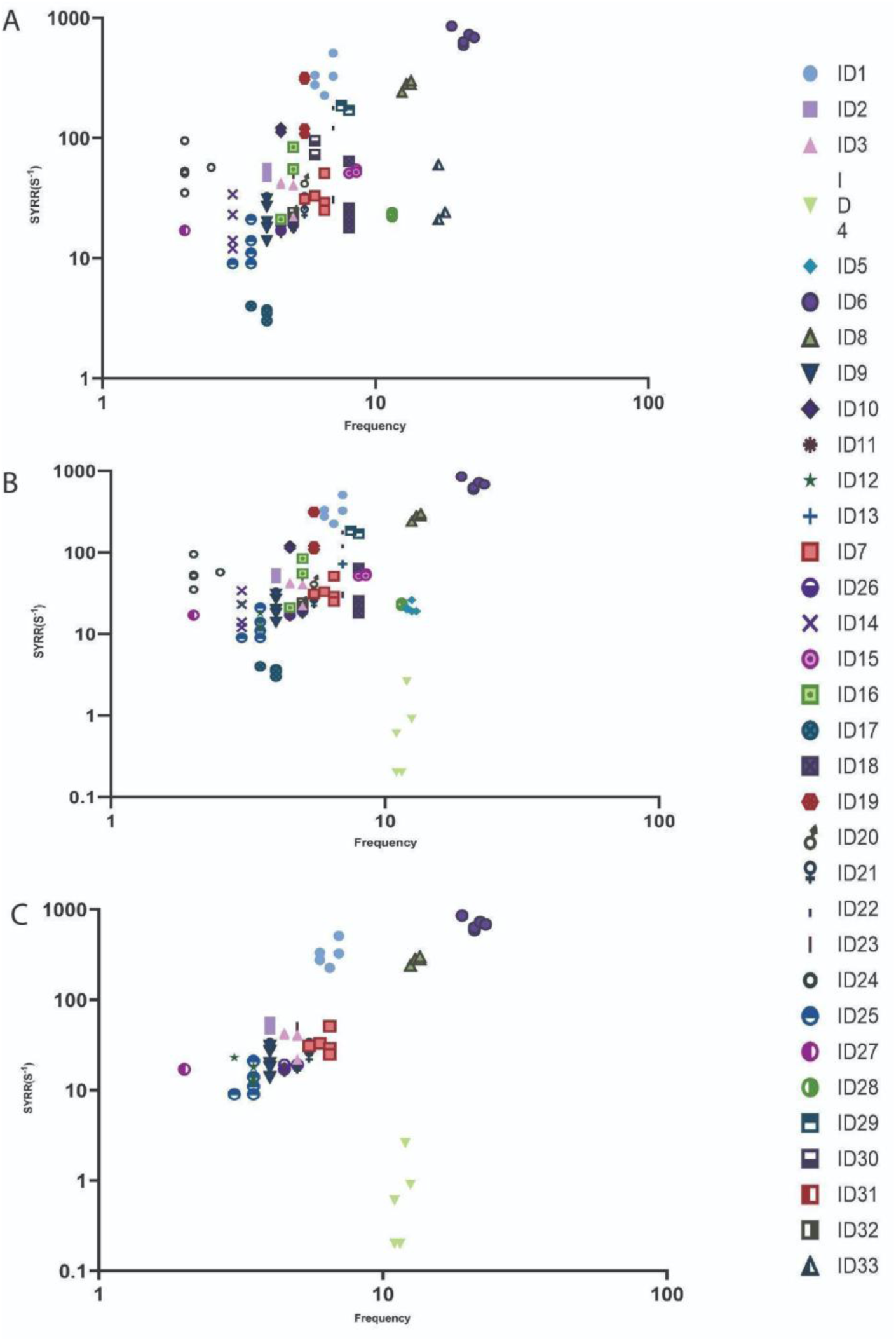
The graphical representation of the distribution of different call types based on log syllable repetition rate and log peak frequency in different seasons. Post-monsoon and pre-monsoon seasons show more varieties of call types than the dry season. A. Pre monsoon B. Post monsoon C. Dry season

This shows a difference in acoustic community assemblage between seasons. The dry season has the least number of call types compared to pre monsoon and post monsoon, but this difference is relatively slight: only ID1, ID6, ID7 and ID 24 are missing in the dry season.

### 5. Acoustic Separation of all call types

The previous analysis shows many call types to be overlapping when separated based on syllable repetition rate and peak frequency. We analysed the acoustic separation of all call types using a hierarchical method where we grouped the callers based not just on their peak frequency and syllable repetition rate but also by incorporating the bandwidth and broader acoustic temporal characteristics as mentioned in Table 1. In addition, we examined clustering using a multivariate principal component analysis using temporal features like syllable repetition rate, syllable period, echeme period and spectral features like peak frequency and bandwidth.

#### 5.a. Acoustic separation in “Triller” call types

Following the PCA, the separation of all the triller call types were visualized relative to the two principal components (Fig. 4.9). Here ID1, ID11, ID15 and ID31 are well separated from the rest of the triller call types. But ID14, ID24 and ID25 show overlaps in the plot. Both syllable types of ID24 also show an overlap in their acoustic characteristics. ID 14 exhibits high individual variation.

The PCA biplot (Fig. 9A) shows that peak frequency and syllable period are negatively correlated to each other and bandwidth and syllable repetition rate are positively correlated. All these four parameters comprise PC1 and PC2.

**Figure 9.**
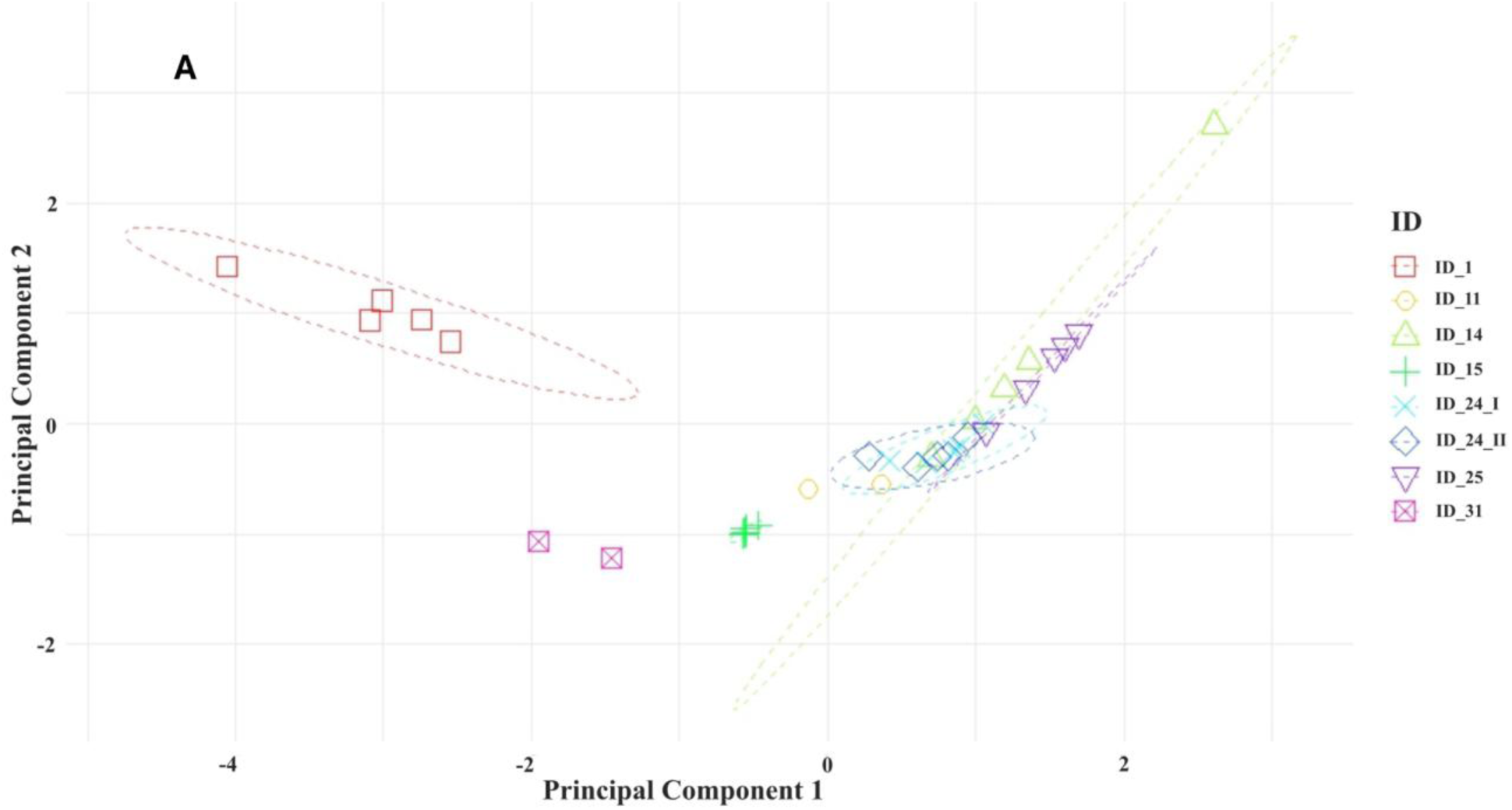
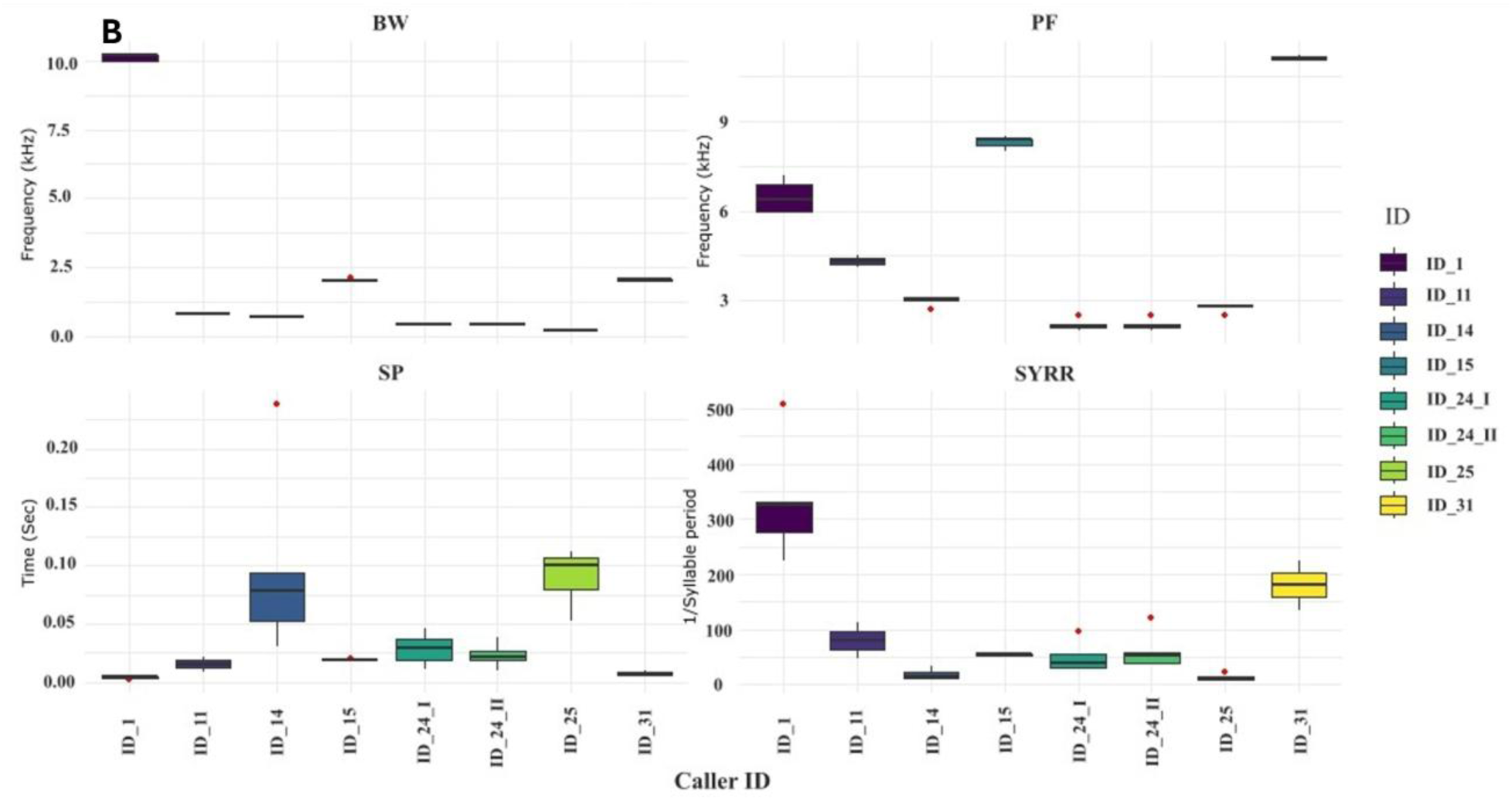
PCA and box plot demonstrating acoustic separation of all Triller call types (A) PCA demonstrating “Triller” call types based on multiple acoustic characters. (B) Biplot showing the acoustic parameter axis directions in each PC. (C) Differences in the bandwidth, peak frequency, syllable period and syllable repetition rate between different triller call types.

When considering all these variables, we find that ID24 and ID25 show differences in syllable structures. ID25 and ID 14 exhibit similarities in syllable structure but there is a slight difference in bandwidth between ID14 and ID 25 (P = 0.05). ID14 and ID24 show differences in frequency bandwidth (P = 0.03).

#### 5.b. Acoustic separation of the high-frequency chirper callers

The separation of high-frequency chirpers with respect to all the temporal and spectral features, including echeme characteristics, was also visualized through dimensionality reduction. It shows a clear separation between the high-frequency narrow bandwidth and broadband chirpers (Fig. 10). The three broad-band high-frequency callers ID6, ID7, and ID8 are well separated from each other and from the narrowband high-frequency chirpers. All different syllable types of ID7 are clustered together. On the other hand, all the narrow-band callers gathered and overlapped.

**Figure 10.**
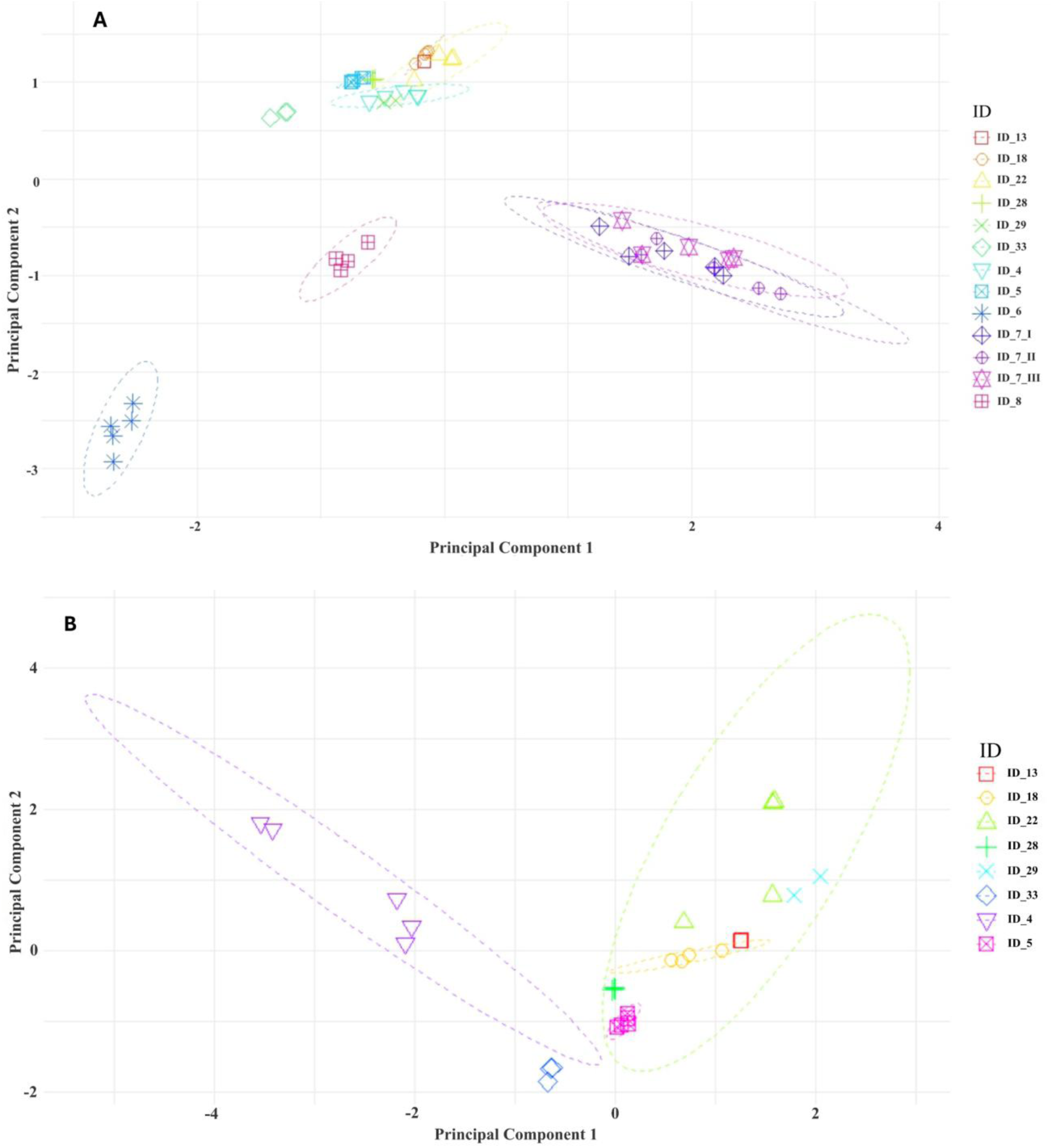
PCA demonstrates clusters of different high-frequency chirper call types based on multiple acoustic characteristics. (A) PCA plot of all high-frequency chirpers; (B) PCA biplot showing the direction of the axes representing different variables (C) PCA plot of only narrow-band high-frequency chirpers.

In the next PCA plot we removed all the broadband callers and plotted only narrow band high frequency callers. In that plot except ID22 all other callers show clear separate clusters. ID22 exhibits inter-individual variation but still separates out from the other callers. It concludes all high-frequency callers are well separated from each other.

#### 5C. Acoustic separation of low-frequency chirpers

A PCA of all the low-frequency chirpers shows that while some calls are well separated from each other, other chirper call types overlap with each other. Chirpers are ID2, ID12, ID24, ID26, and ID30 form well-separated clusters. But ID3, ID9, ID16, ID23, and ID32 show considerable mutual acoustic overlap. ID19 shows partial overlap with ID21. ID27 and ID17 exhibit high levels of variation between individuals (Fig. 11).

**Figure 11.**
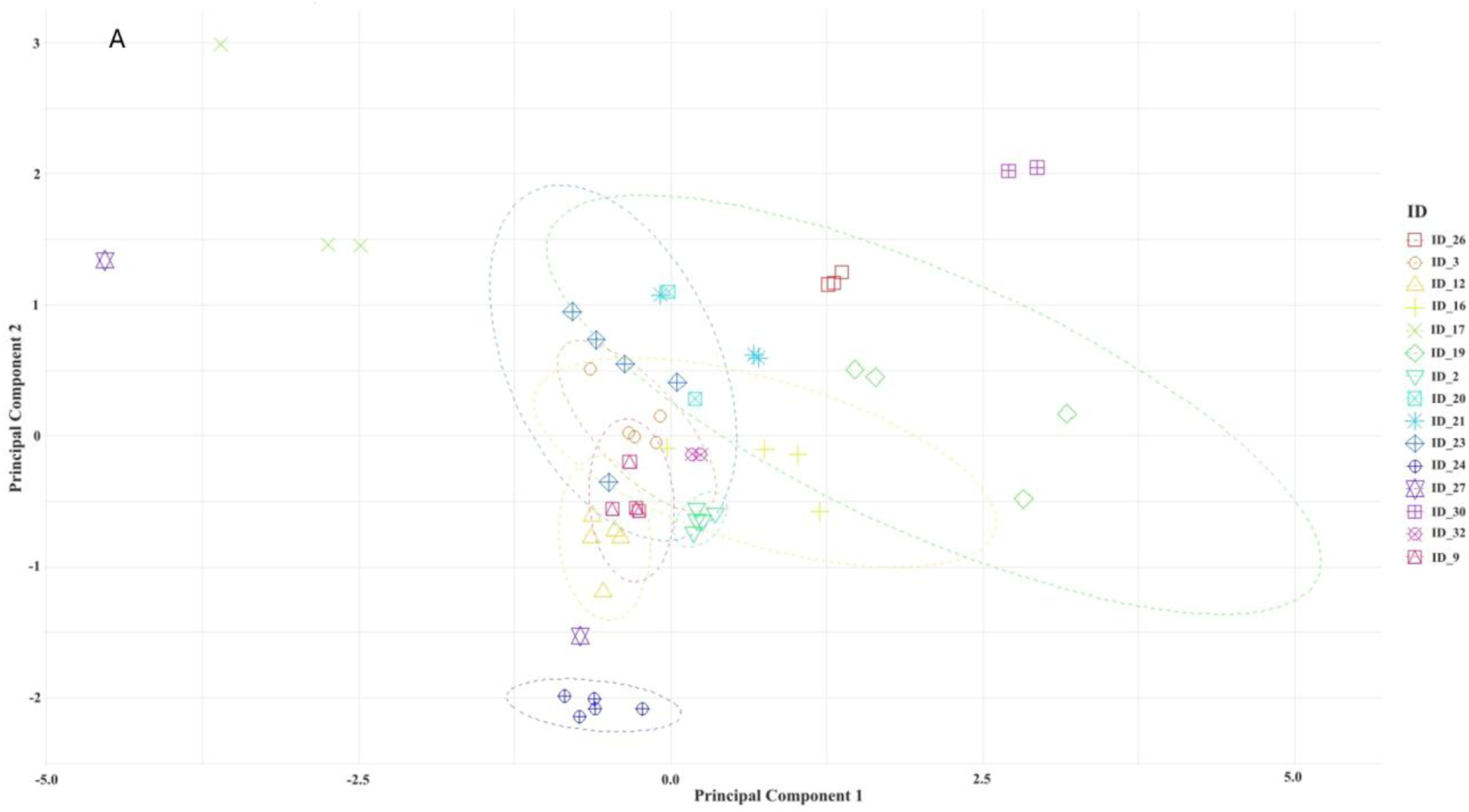
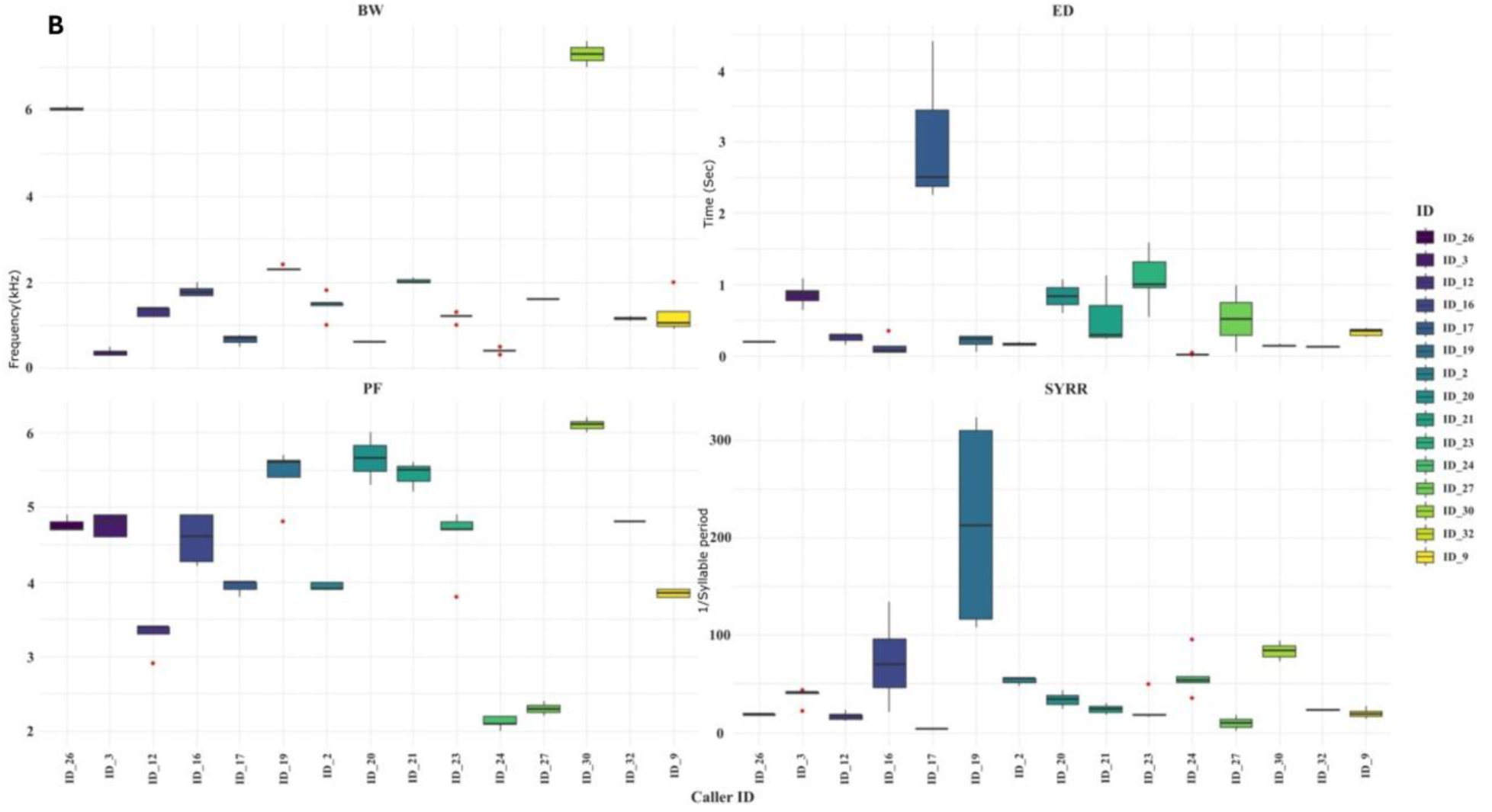
PCA demonstrates clustering of different low-frequency chirper call types based on multiple acoustic characters. A. and B.

The PCA biplot shows peak frequency, bandwidth, and syllable period as the parameters contributing the most to cluster separation. We separately examined variation in each parameter with respect to different low-frequency chirper call types. ID9 is significantly different in bandwidth from ID16 and ID 3 (P = 0.02 and 0) and significantly different in peak frequency from ID23 and ID32 (P = 0.008 and 0.007 respectively), although it shows no significant difference in syllable structures with them. ID3 is significantly different in bandwidth from ID16, ID23, and ID32 (P = 0, 0.001, 0.002 respectively). ID16 is significantly different in terms of echeme period from ID23 (P = 0.01).

### 6. Acoustic space use (ASU) analysis

This map illustrates the seasonal distribution of frequencies in both transects, with darker colors indicating a higher proportion of callers in that frequency bin present at a site (Fig. 12). Darker shades suggest that a larger number of call types within the site produce calls at those frequencies, which can be interpreted as regions where signal masking is more likely to occur due to overlapping frequency ranges. This representation identifies the presence/absence of each call type in each transect (summed across the ten sites within each transect) in each frequency band, and therefore this measure does not count the number of callers from each call type within those frequency bins.

**Figure 12.**
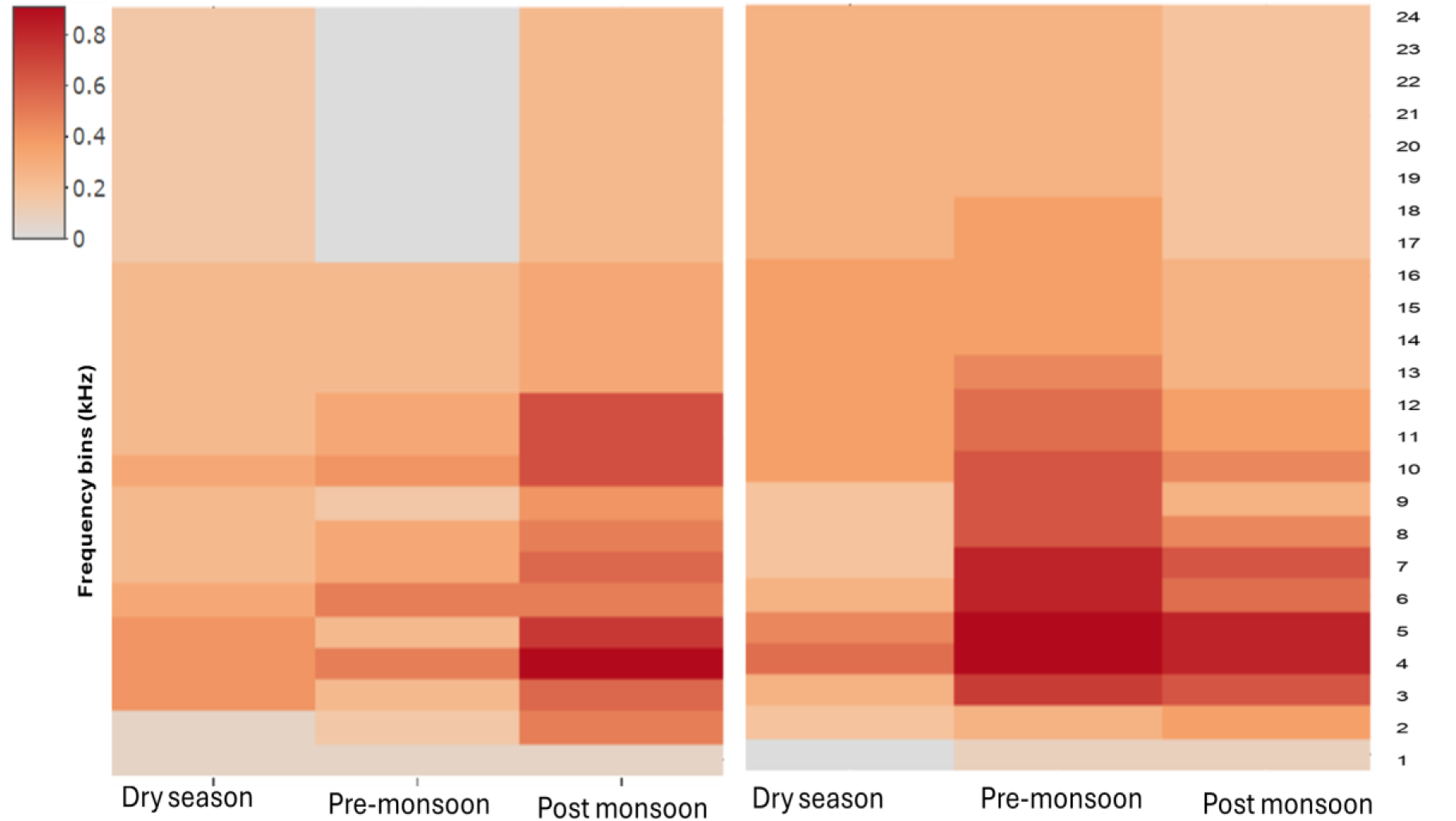
Heat Map of acoustic space use (ASU) by different callers in different seasons. The plot on the left side shows transect1 and the plot on the right side depicts transect 2. Each bin on the y-axis represents a 1 kHz frequency bin for a particular season. The color scale of the heat map is on the left.

In Transect 1, the post-monsoon season exhibits the broadest frequency coverage compared to the pre-monsoon and dry seasons. Within the post-monsoon season, there is a concentration of callers in the 4–5 kHz and 10–12 kHz ranges.

In Transect 2, more of the spectral acoustic space is utilized across seasons. However, the distribution varies. In Pre-monsoon most callers are concentrated within the 3–12 kHz range. In the post-monsoon season, the range shrinks - most activity occurs in the 3–7 kHz range. During the dry season callers are predominantly found at 4 kHz.

The proportion of acoustic space use (ASU) varies across months and between the two transects, indicating seasonal and local spatial differences (Fig. 13). In transect 1, the post-monsoon season exhibits the highest ASU proportion, accounting for approximately 40%, followed by 26% in the dry season and 22% in the pre-monsoon season. In transect 2, the pattern differs, with the pre-monsoon season showing the highest ASU proportion at 46%, followed by 35% in the post-monsoon and 29% in the dry season. These findings demonstrate that ASU proportions are not consistent across seasons or even transects that are near one another and were chosen as replicates, highlighting variations in the temporal and microhabitat related dynamics of acoustic activity. These patterns provide insight into seasonal variations in frequency use and potential signal masking within the soundscape.

**Figure 13.**
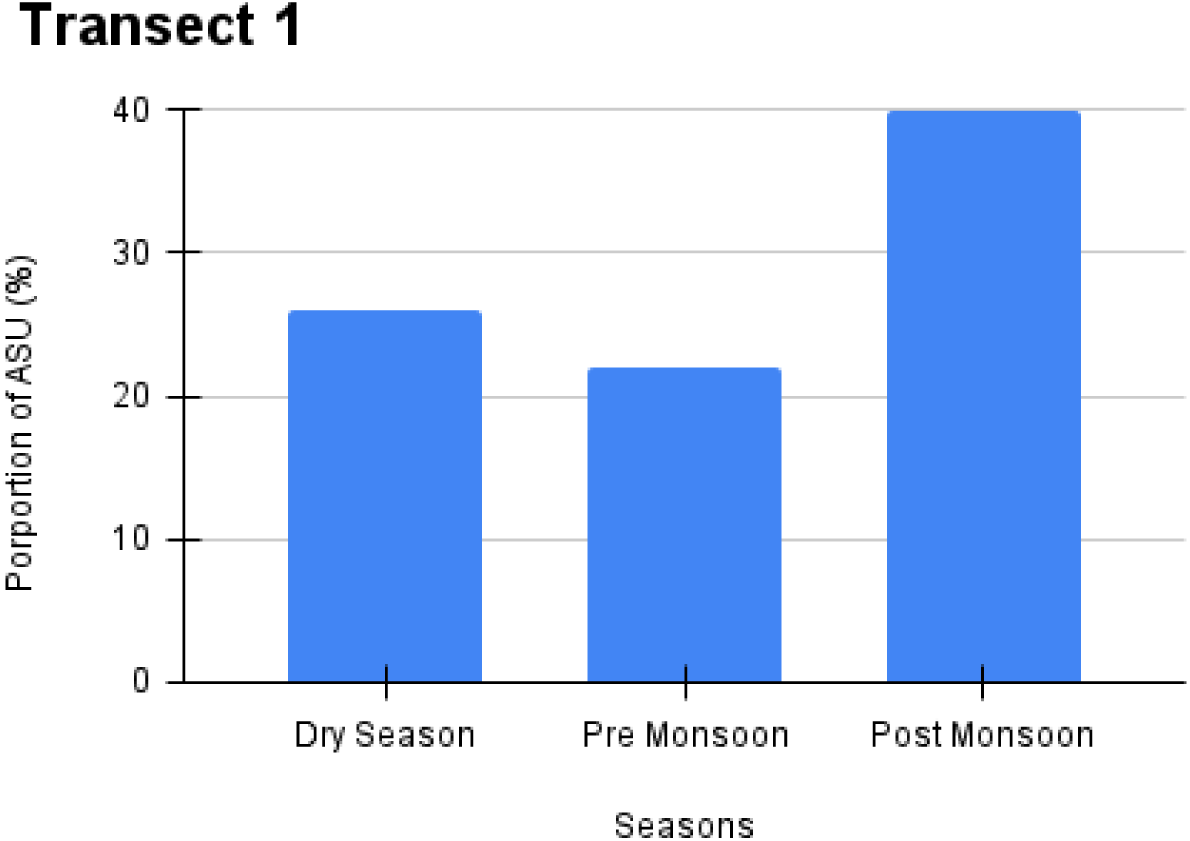
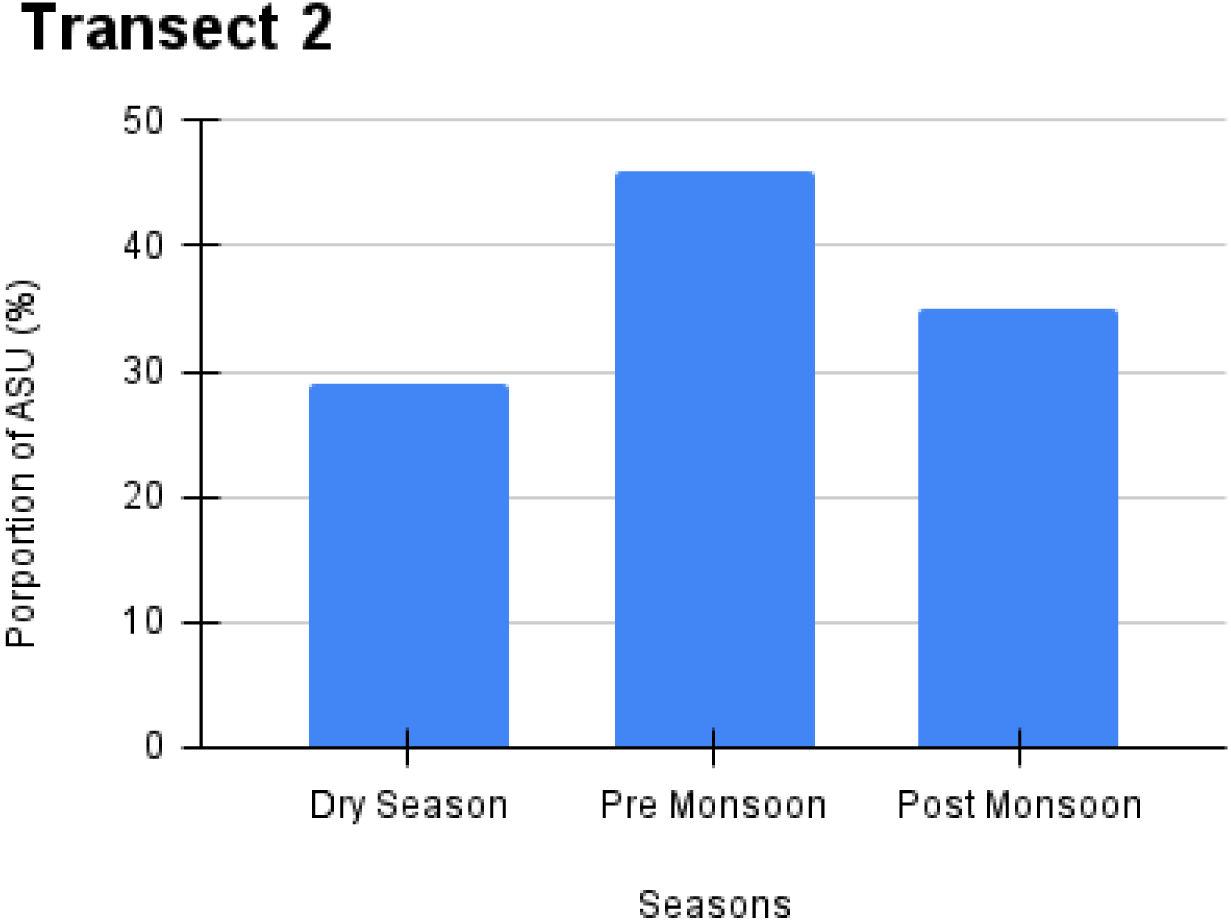
The difference in acoustic space use (ASU) proportion in each season. (top) Transect site 1 and (bottom) Transect site 2.

Across both transects and seasons, there are very few callers producing calls in the higher frequency range of 17–24 kHz. This highlights the dominance of low and mid-range frequency callers relative to the human audio range in the acoustic environment, consistent with a strong presence of ground crickets (family Gryllidae) and suggests limited use of higher frequency bands by bushcrickets (family Tettigoniidae) species in this area.

### 7. Occurrence of different call types throughout all seasons

Heat maps were created with the occurrence rates of each call type in each season (Fig. 14). This exhibits variation in the occurrence of particular call types in each season. There are some call types that are present only in one season or even only in one transect, like ID33 and ID32. Multiple call types are present throughout the seasons. Out of that, some call types here have high occurrence rates in every season; one such example is ID6, which we had identified as *Ducetia sp* (Tiiwari and Diwakar, 2022). ID7 shows a high occurrence only during the post-monsoon and dry seasons.

**Figure 14.**
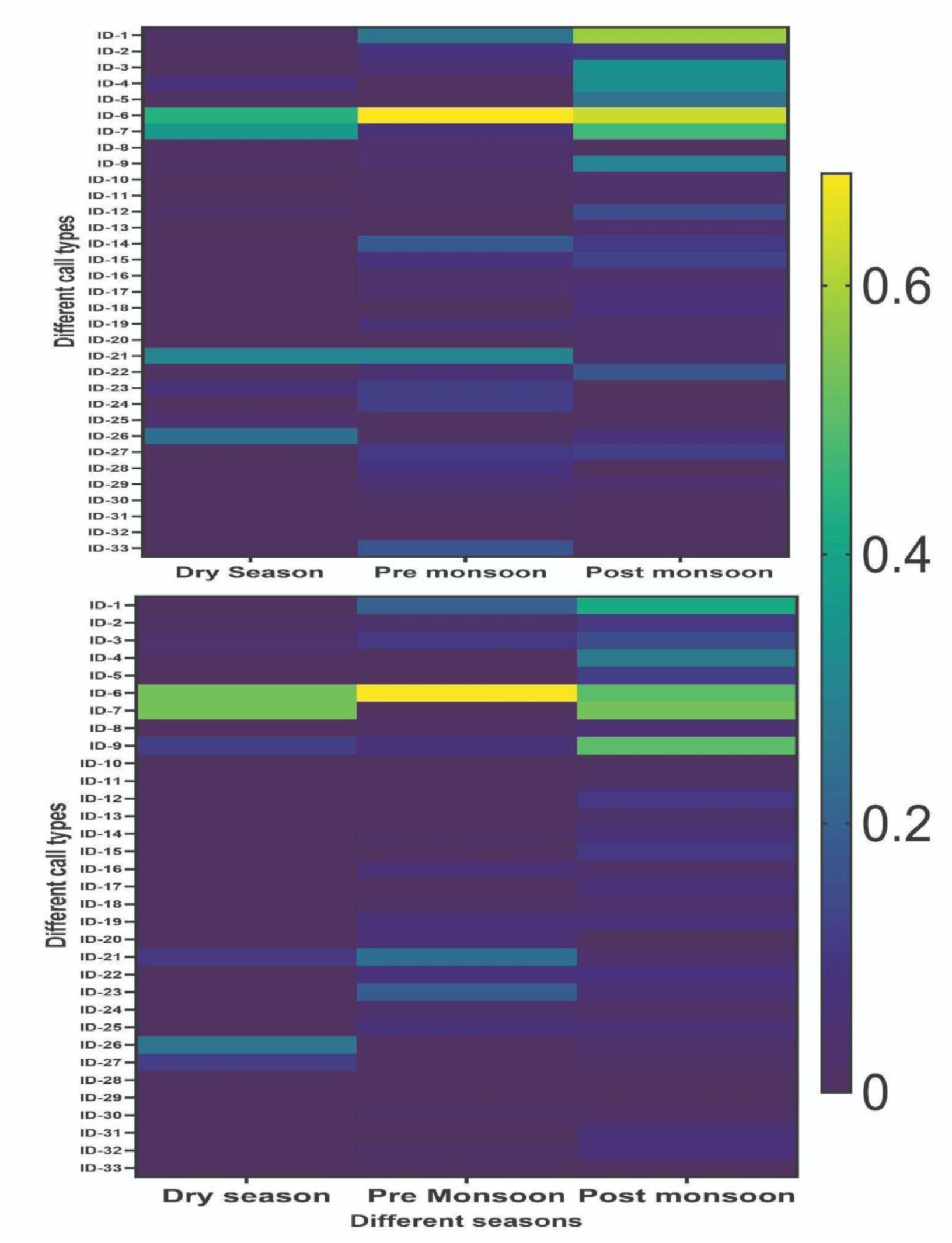
Heat map of call type occurrence for each call type in different seasons at two transect sites. Color gradients denote the occurrence of particular call types, with lighter colors representing higher occurrence.

The occurrence rate was calculated with the following formula:

**Occurrence Rate** = (Number of segments where the caller is detected) / (Total number of segments the season)

This indicates the presence and absence of each call type in different seasons at different sites. The color bar indicates the coding of the occurrence of the call type that gets recorded by the device at each of the two transects in each season.

The occurrence rate of each call type across the seasons shows considerable variation. In the dry season most of the callers have lower occurrence or even complete absence compared to post-monsoon or pre-monsoon. We also test for significant differences in call type occurrence across seasons, using a Kruskal-Wallis test, considering only the call types present at some level across all three seasons. In transect 1, ID3, ID4 and ID9 show a significant difference between post-monsoon and the other two seasons (p = 0.02, 0.001, 0.05 respectively) but ID8 alone shows higher pre-monsoon and dry season occurrence than post-monsoon (p = 0.01). In transect 2, ID4, ID9, and ID1 have significant differences between post-monsoon and the other two seasons (p = 0.03, 0.005, 0.001 respectively), but other call types do not show a significant difference in occurrence rates.

#### Transect 1

A linear mixed-effects model was used to analyze the effect of season on occurrence values, with call type ID included as a random effect to account for variability across individual call types (Fig. 15A). The model revealed a significant effect of the post-monsoon season compared to the dry season (estimate = 0.078, SE = 0.022, t = 3.56, p < 0.001). The pre-monsoon season showed a positive but non-significant effect relative to the dry season (estimate = 0.039, SE = 0.023, t = 1.72, p = 0.087). These results suggest that occurrence values were higher in the post-monsoon season than in the dry season, with the pre-monsoon season showing a non-significant increase compared to the dry season.

**Figure 15.**
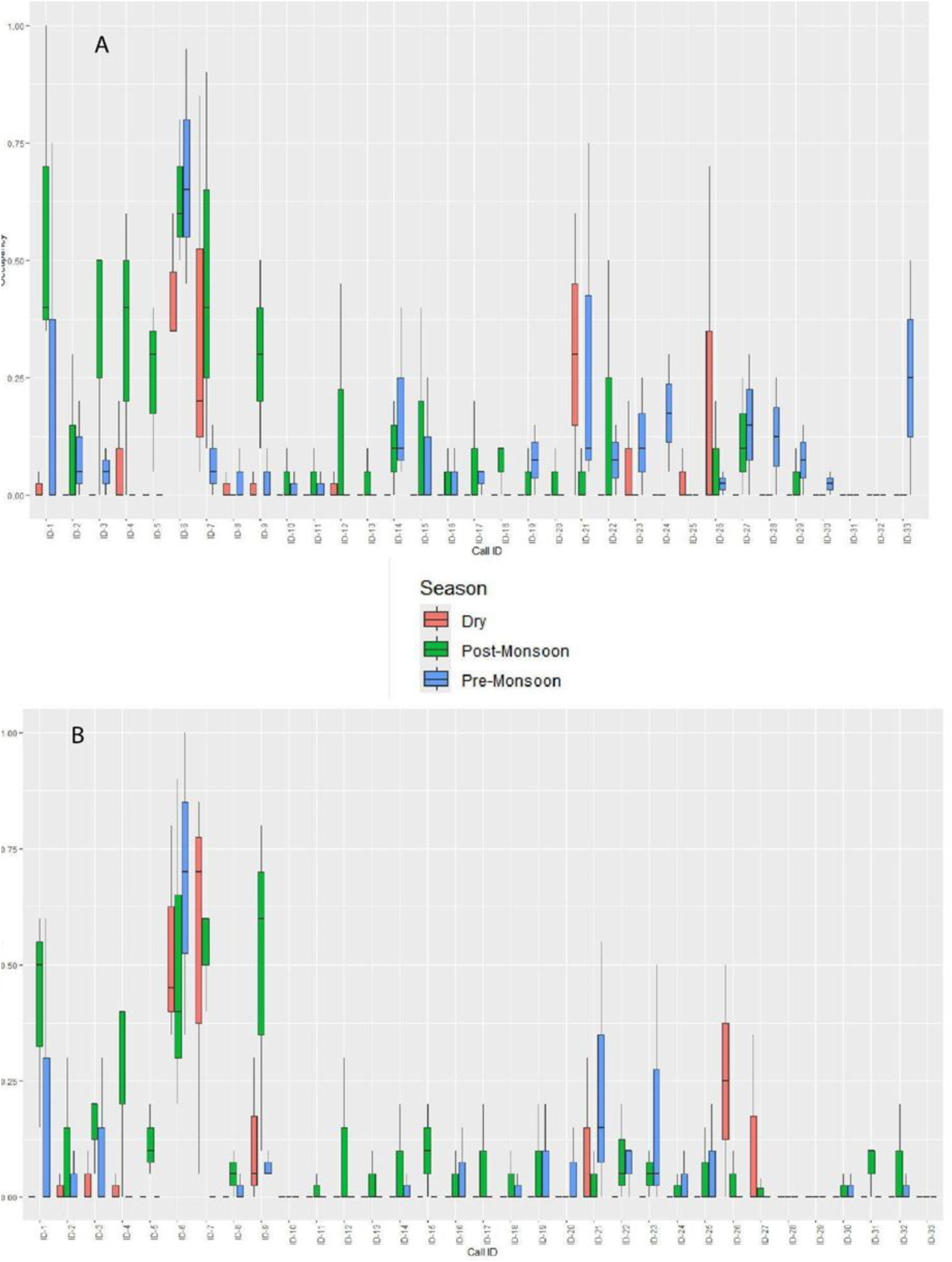
Call type occurrence showing variation between seasons. (A) Transect site 1 and (B) Transect site 2.

**Figure 16.**
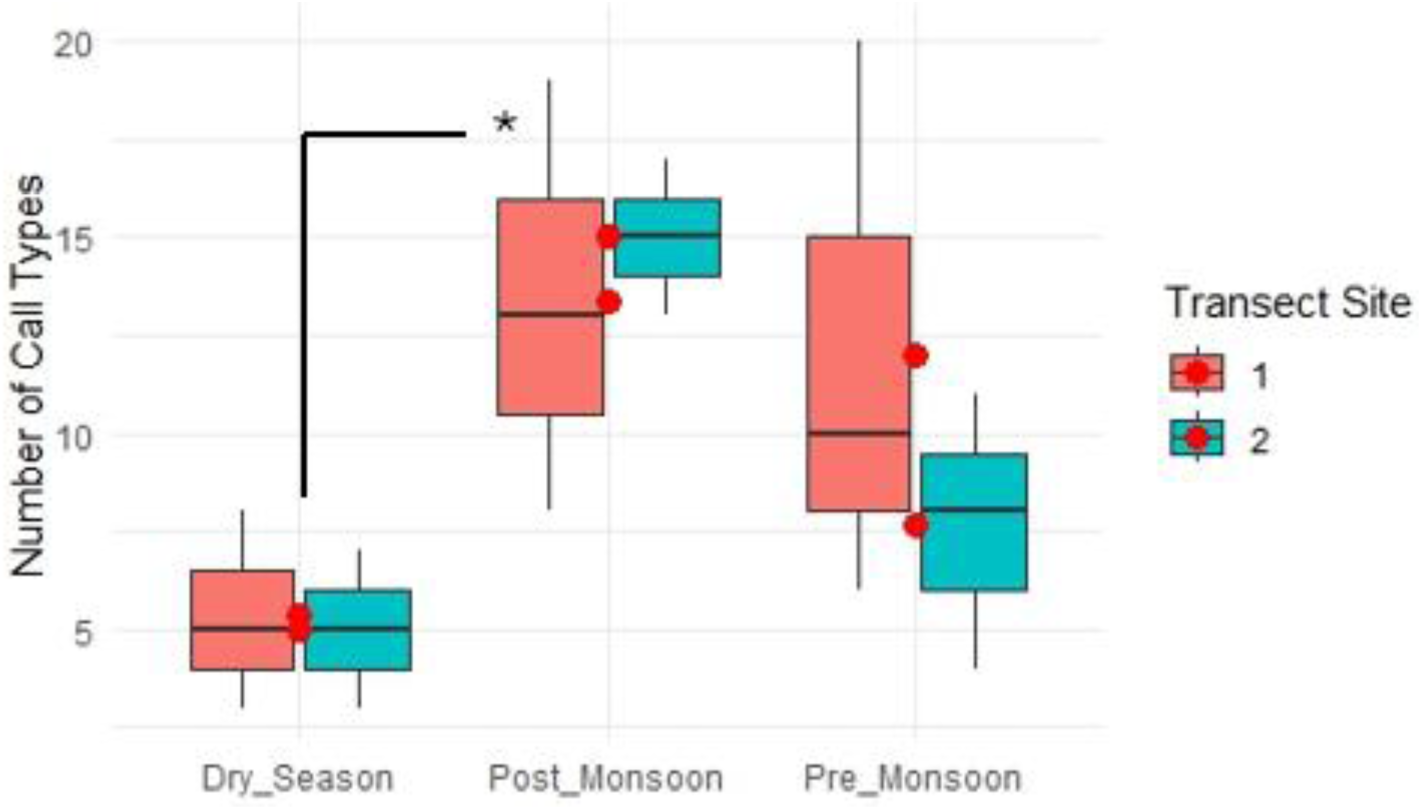
The number of call types present in each season.

#### Transect 2

A linear mixed-effects model was conducted to assess the effect of season on occurrence values, with call types included as a random effect to account for variation across call types (Fig. 15B). The random intercept for ID showed a variance of 0.0118 (SD = 0.109), indicating moderate variability in occurrence across different call type IDs. The residual variance was estimated at 0.0171 (SD = 0.131). For fixed effects, there was a significant effect of the post-monsoon season compared to the dry season (estimate = 0.0565, SE = 0.0186, t = 3.04, p = 0.0026). This suggests that occurrence rates were significantly higher during the post-monsoon season. The pre-monsoon season, however, did not show a significant effect on occurrence compared to the dry season (estimate = 0.0061, SE = 0.0186, t = 0.33, p = 0.744). Model diagnostics indicated a good fit, with a REML criterion at convergence of -285.4.

### 8. Seasonal variation in call type composition

The cumulative data on the number of call types in each season shows variations within each season due to monthly variations in the number of call types within each season. Still, the dry season has a significantly lower call number throughout all months in both transects compared to post-monsoon and pre-monsoon seasons (p = 0.003).

Not just the number of call types, but also the call type composition differs in each month within the season. The variation in call type composition in each season can be visualized using NMDS for each transect site.

#### Transect 1

The statistical analysis for transect site 1 revealed no significant seasonal differences in the diversity and call type composition (Fig. 17A). A PARANOVA indicated no overall significant seasonal effect (F = 2.99, p = 0.126), and post-hoc Tukey tests showed no significant pairwise differences between seasons (all p > 0.1). The par ANOVA on NMDS Axis 1 scores did not reveal a significant effect of season (F = 4.35, p = 0.13), and NMDS Axis 2 scores also showed no significant variation across seasons (F = 0.47, p = 0.666). These findings suggest that seasonal changes do not significantly influence call type composition at this site.

**Figure 17.**
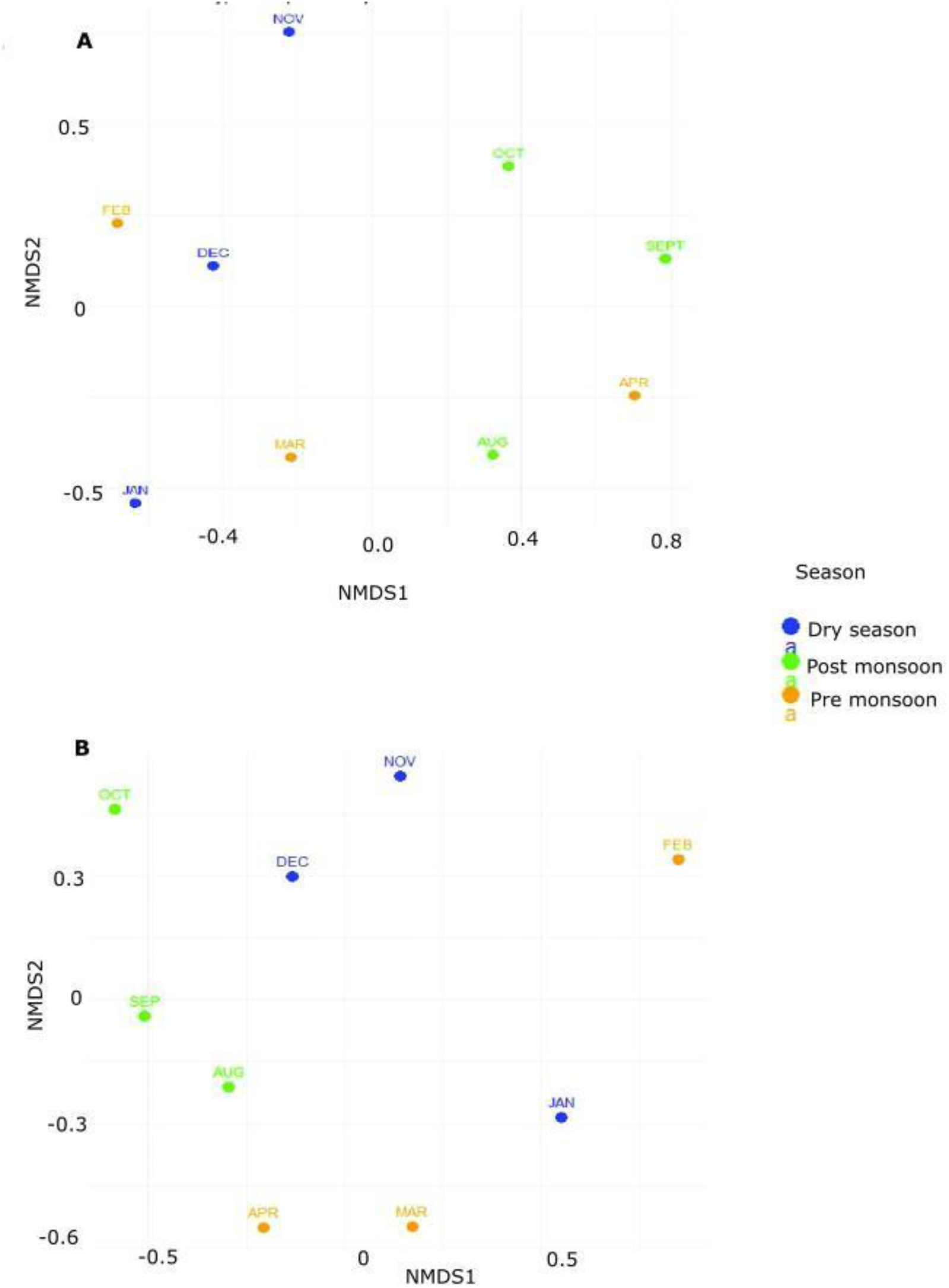
Non-parametric multivariate analysis of the composition of different call types in different seasons at both sites (A: transect site 1, B: transect site 2). Each dot represents one month from each season. The monsoon could not be represented due to the masking effect of the noise of the rain over and above any acoustic sounds.

**Figure 18.**
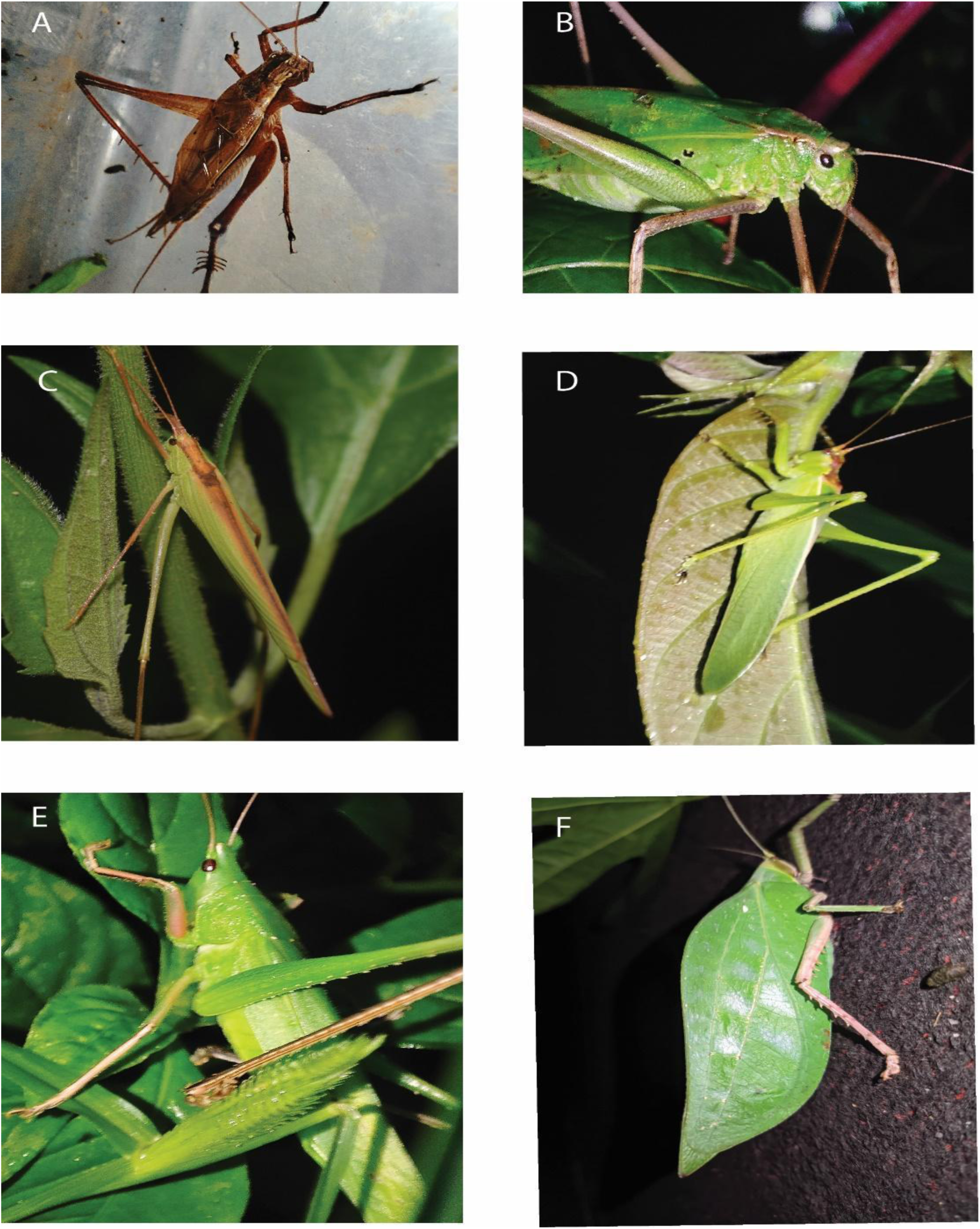
Photographs of a few Ensiferans from Nongkhyllem WLS. A. *Gymnogryllus sp* (ID-9), B. *Mecopoda “Complex”* (ID-7), C. *Ducetia assamica* (ID-6), D. *Hexacentrus unicolor* (ID-8), E. Conocephalinae and F. Pseudophyllinae.

#### Transect 2

The analysis for transect site 2 demonstrated significant seasonal differences in call type composition but no significant differences in NMDS axes, indicating variability in diversity but not in overall call type composition. PARANOVA showed a significant seasonal effect (F = 6.50, p = 0.0315), with Tukey post-hoc tests revealing that post-monsoon had significantly higher diversity compared to the dry season (p = 0.0299) (Fig. 17B).

## Discussion

This passive acoustic monitoring study identified 33 distinct call types over one year, excluding the monsoon period. This research represents the first bioacoustic study of Ensiferan insects in Meghalaya, one of the least explored states in Northeastern India. The study relied entirely on passive acoustic recordings, and as a result, the specific identity of the callers remains unknown. All recorded calls are presumed to belong to nocturnal species, as the recordings were conducted exclusively at night.

The focus of this study was Ensiferan callers, a group that includes crickets bushcrickets, which typically produce highly distinctive calls. Ensiferan calls are characterized by syllables generated by single-wing strikes. These syllables may occur in grouped patterns, forming chirps, or as continuous non-grouped syllables, forming trills. These features make Ensiferan calls acoustically distinct both to the ear and in oscillograms, distinguishing them from the more complex calls of nocturnal birds. Additionally, bat calls differ markedly due to their use of frequency-modulated ultrasonic pulses. The study’s 48 kHz sampling rate was insufficient to capture high-frequency bat echolocation calls, thereby precluding confusion with such signals. We were able to use focal recordings with ultrasound recorders with specific broadband callers whose range appeared to extend beyond 24 kHz, like *Hexacentrus* (ID8), *Ducetia* (ID6) and *Mecopoda* (ID7). We may well have missed narrow band callers whose fundamental frequency is above 24 kHz using the Passive Acoustic Monitoring method.

The primary acoustic similarity could occur with anuran calls, as certain frog species, such as cricket frogs, mimic cricket-like sounds. However, the study’s sampling rate also excluded the detection of any potential high-frequency Ensiferan callers (with a fundamental frequency higher than 22 kHz) that might exist in the region. Also, as the device was placed near the forest understory, it missed any ensiferan callers with high-frequency, low-intensity callers as high-frequency calls travel less far, and the device only enabled to detect the sound in its detectability range.

In the field, psychoacoustically distinguishing all call types was challenging. However, using spectral FFT views, it became evident that all calls were acoustically distinct from one another. Taxonomic identification was possible for only four species, each from different genera, which were abundant and exhibited loud, broad-range calls: *Hexacentrus unicolor, Gymnogryllus sp., Mecopoda elongata “Complex,”* and *Ducetia assamica*. Notably, *Ducetia assamica* was recently described by Tiwari and Diwakar (2023) in Assam, while the *Mecopoda elongata “Complex”* is reported here for the first time from the Garo Hills of Meghalaya (Ghosh et al., 2025).

The number of call types varied seasonally and spatially across the transects. In Transect 2, call types ID10, ID28, ID29, and ID33 were absent, while ID31 and ID32 were absent in Transect 1. The acoustic community exhibited a composition dominated by low-frequency callers (<4 kHz) and fewer high-frequency callers (>10 kHz) with high syllable repetition rates. Most call types were concentrated in the 4–8 kHz frequency range, exhibiting diverse syllable repetition rates. Seasonal differences were evident: the dry season had fewer call types, with a more conserved acoustic community predominantly in the 4–8 kHz range with lower syllable repetition rates. In contrast, the pre-monsoon and post-monsoon seasons exhibited a broader frequency range and more varied syllable repetition rates.

As the individual call types were extracted from passive recording, many low-intensity callers got partially masked by the high-intensity broad-band callers or noise. Therefore, it was difficult to have enough samples to perform temporal analysis for a few call types that were only occasionally represented in the dataset. This constitutes a general limitation of passive recordings.

### Acoustic separation

Multivariate analysis using PCA revealed distinct separation of triller call types, except for ID14, ID25, and ID24, which overlapped on broader scales but were distinguishable on finer temporal and spectral scales. For high-frequency chirper call types, high-frequency broadband calls formed separate clusters. Narrowband high-frequency calls showed clear separation at finer resolutions. Among low-frequency callers, ID3, ID9, ID16, ID23, and ID32 exhibited overlapping acoustic features along the PC1 and PC2 axes. However, we found subtle separations based on minute spectral differences (∼500 Hz) and echeme period, distinguishing these call types on a finer scale.

### Acoustic space use

In the periphery of the Nongkhyllem Forest Reserve, the acoustic environment demonstrates engagement across the 1–16 kHz frequency range throughout the year at both sites. During the post-monsoon and pre-monsoon seasons, the spectrum extends further, with full engagement observed from 1-24 kHz. However, in Site 2, the entire spectrum is consistently occupied regardless of the season. The most utilized acoustic space is between 3–7 kHz at both sites across all seasons, indicating that most call types fall within this frequency range. The proportion of acoustic space usage varies with season and location. In the dry season, both sites exhibit lower acoustic space use. At Site 1, post-monsoon shows the higher proportion of acoustic space usage, followed by pre-monsoon, while at Site 2, pre-monsoon has a higher proportion of usage than post-monsoon in one year of data.

### Occurrence of call types

The occurrence rate of call types also varies seasonally at each site. Among the call types, ID7 exhibited the highest occurrence rate throughout the year. Occurrence rates differed significantly between the post-monsoon season and the other two seasons, with post-monsoon showing the highest overall occurrence. However, there was no significant difference in occurrence rates between the dry and pre-monsoon seasons at either site. The monthly composition of call types within each season varied, leading to some seasonal differences in call type composition. However, non-metric multivariate analysis indicated no overall significant differences in call type composition between seasons across both sites. This suggests that while there are seasonal differences in calling activity, the contribution of individual call types to the acoustic community remains relatively consistent between seasons.

### Global Ensiferan diversity

Tropical and subtropical forests are among the most biodiverse regions globally in terms of species diversity and resource richness. India hosts four biodiversity hotspots, all of which belong to tropical or subtropical forest ecosystems. Our study site falls within the Indo-Burma biodiversity hotspot. Numerous studies have investigated the species diversity of Ensiferans (crickets and bushcrickets) in tropical forests in India and worldwide.The pioneering study by Diwakar and Balakrishnan (2007) reported the species and acoustic diversity of Ensiferans in the tropical forests near Kudremukh National Park, part of the Western Ghats biodiversity hotspot in South India. They documented 20 Ensiferan species from the Gryllidae and Tettigoniidae families. Their findings revealed variations in temporal and spectral acoustic structures among species. Although the study did not report seasonal variation, it provided insights into the diel activity of Ensiferans. They observed most ground crickets producing calls in the 3–7 kHz range, with syllable structures varying from simple chirps to trills. Bushcrickets calls included both high- and low-frequency callers. Subsequent studies expanded on these findings (Jain & Balakrishnan., 2010). Tiwari and Diwakar (2022) investigated bushcrickets diversity in Assam (Hoolock Gibbon Wildlife Sanctuary), also from the Indo-Burma hotspot, and Goa (Bhagwan Mahaveer Wildlife Sanctuary) from the Western Ghats biodiversity hotspot. They reported 15 bushcrickets species and documented seasonal variation in bushcricket assemblages between the post-monsoon and dry seasons. For instance, *Hexacentrus* species, *Mecopoda “Helicopter”*, and *Mecopoda “Triller”* were recorded exclusively in the post-monsoon season, whereas *Mecopoda fallax* and *Ruspolia indica* were restricted to the dry season. Conversely, *Ducetia* species were observed year-round. In our study also, we found *Ducetia sp.* (ID6) to be consistent throughout the year and *Mecopoda “Complex,”* found in the dry season and pre-monsoon compared to the post-monsoon season. One possible reason could be that *Mecopoda* is one of the largest bushcrickets in the understory, producing the loudest broadband call with a high duty cycle which is well represented in recordings in comparison to calls that are softer, or have a low duty cycle. It is worth noting, however, that Jain et al. (2014) show that *Mecopoda* does not produce a masking interference problem for other callers in the Western Ghats. On the other hand *Ducetia sp* produces low intensity chirps, but other ecological factors might be responsible for its year-round presence.

Globally, studies in paleotropical and neotropical regions have revealed significant variations in Ensiferan assemblages. In Indonesia’s tropical forests, Syahlan et al. (2017) reported 24 Ensiferan species, although their study lacked acoustic data. Earlier, Riede (1997) conducted research in the Borneo rainforests, highlighting a rich diversity of Ensiferans and considerable spatiotemporal separation in their acoustic niches compared to subtropical and temperate forests.

In the Neotropics, studies in Ecuador and Panama have further enriched our understanding. Nischk and Riede (2015) found that acoustic assemblages in Ecuador’s cloud forests decreased in diversity with increasing altitude, with a total of 35 Ensiferan species recorded. In temperate regions, Roca and Proulx (2016) reported 15 Ensiferan species in Canada. Their findings demonstrated a positive relationship between acoustic heterogeneity and species richness, emphasizing the usefulness of acoustic diversity as an indicator of biodiversity.

The acoustic niche partitioning hypothesis posits that call types evolve to separate out in overall acoustic space, as well as in terms of the spatial location and time of calling, to reduce masking interference. Schmidt et al. (2012) described greater spectral niche partitioning of Ensiferan species than expected by chance, with the size of the acoustic niche increasing with spectral distance to the nearest neighbor. These studies collectively highlight the vast acoustic and species diversity of Ensiferans across tropical and subtropical ecosystems, contributing valuable insights into their ecological and acoustic dynamics.

Most previous studies on Ensifera relied on psychoacoustic methods with active recording techniques to collect data. However, passive acoustic monitoring (PAM) has emerged as a valuable tool in ecological studies of vocalizing organisms. The first application of PAM for bushcrickets was recently conducted by Symes et al. (2022) in the tropical forests of Panama. They passively recorded forest sounds and compared their findings to a pre-existing acoustic and taxonomic database developed by ter Hofstede (2020). By integrating this database with machine learning, they were able to identify and categorize all the recorded calls efficiently.

In our study, we recorded a comparatively higher number of call types than other paleotropical forests, potentially reflecting the diversity of Ensiferan species in this region. However, since this study was conducted in a new geographical area without an established database of Ensiferan calls, we were unable to actively use machine learning for automatic call identification. This highlights the need for further research and database development in such unexplored ecosystems. PAM has been increasingly applied across diverse taxa including cicadas, birds, aquatic animals, primates, and anurans (Clink et al., 2020; Ross et al., 2021; Wood et al., 2023; Hill et al., 2018; Ramesh et al., 2023; Desjonquers et al., 2019).

Many of these studies have explored acoustic space use (ASU), which quantifies the spectral space occupied by specific taxa (Aide et al., 2017; Ramesh et al., 2023). Such analyses provide insights into ecosystem health by allowing long-term comparisons across habitat types. Ensiferans, for instance, can serve as bioindicators due to their sensitivity to environmental changes. Shifts in their acoustic behavior or population dynamics can indicate habitat alterations, including deforestation, land-use changes, or anthropogenic disturbances (Shieh et al., 2012). Reduced vegetation density also affects the transmission of sound waves, altering the quality and range of insect calls (Riede, 1993).

Similar to bioacoustic or acoustic indices, we calculated acoustic space use proportion to understand the diversity of ensiferan sounds throughout seasons. The bioacoustic index is usually focused only on overall biological sounds and audible range frequencies (Boelman, et al., 2007). But since we are interested in a specific taxon (Ensifera), which spans audible to ultrasound frequency ranges, it is difficult to calculate acoustic indices that measure the presence of Ensiferan callers without a full Ensiferan call reference database.

This study highlights the diversity of Ensiferan call types and seasonal variation in acoustic space use within the sub-tropical forests of the Indo-Burma biodiversity hotspot, specifically the Nongkhyllem Wildlife Sanctuary. By leveraging passive acoustic monitoring (PAM), it provides valuable insights into how environmental factors correlate with acoustic diversity in a non-invasive manner (Sugai et al., 2019). The findings underscore PAM’s potential as a model approach for studying Ensiferan diversity, offering foundational data for quantifying call diversity (Gerhardt & Huber, 2002; Greenfield, 2002).

Future work aims to establish a comprehensive call database with taxonomic identification of all Ensiferan species in Nongkhyllem. This database will serve as a training dataset for machine learning algorithms to classify calls to the genus or species level, addressing limitations in species identification. Such advancements will facilitate studies on fine-scale niche partitioning and enable the use of acoustic indicators to monitor ecosystem health, supporting conservation in the face of deforestation and climate change (Riede & Balakrishnan, 2024).

## Acknowledgments

We thank the State Forest Departments of Meghalaya for permission to conduct fieldwork in protected areas. We are grateful to Dr. Shivani Krishna, and Dr. Manjari Jain, for academic guidance. We are thankful to Dr. Meghna Agarwala for helping us with the shape files for QGIS. We also extend gratitude to our field guide Mr. Iniyas and Mr Overson Lyngdoh for their contribution and help in field collection. We acknowledge the immense support from the Khasi community people from Umling village of Ri-Bhoi district, Meghalaya during our field stay. I would like to mention Dr. Sushweta Mahalonbish and Vivek Dasoju for helping in part of the field collection and Ashique Rahman for establishing the field station. We also extend our gratitude to our colleagues at the Biology, Psychology, and Environmental Studies departments at Ashoka University.

## Authors contribution

Conceptualization and idea AG & BKR, Field site selection AG, RW & BKR, Field station establishment AG RW & BKR, Field data collection AG & JB, Data analysis and statistics AG & JB, Writing AG, editing BKR, Supervision and funding BKR. All authors have read and agreed to the published version of the manuscript.

