## Supplementary data for "Passive acoustic monitoring of Ensiferan calling diversity in a sub-tropical forest of Northeast India"

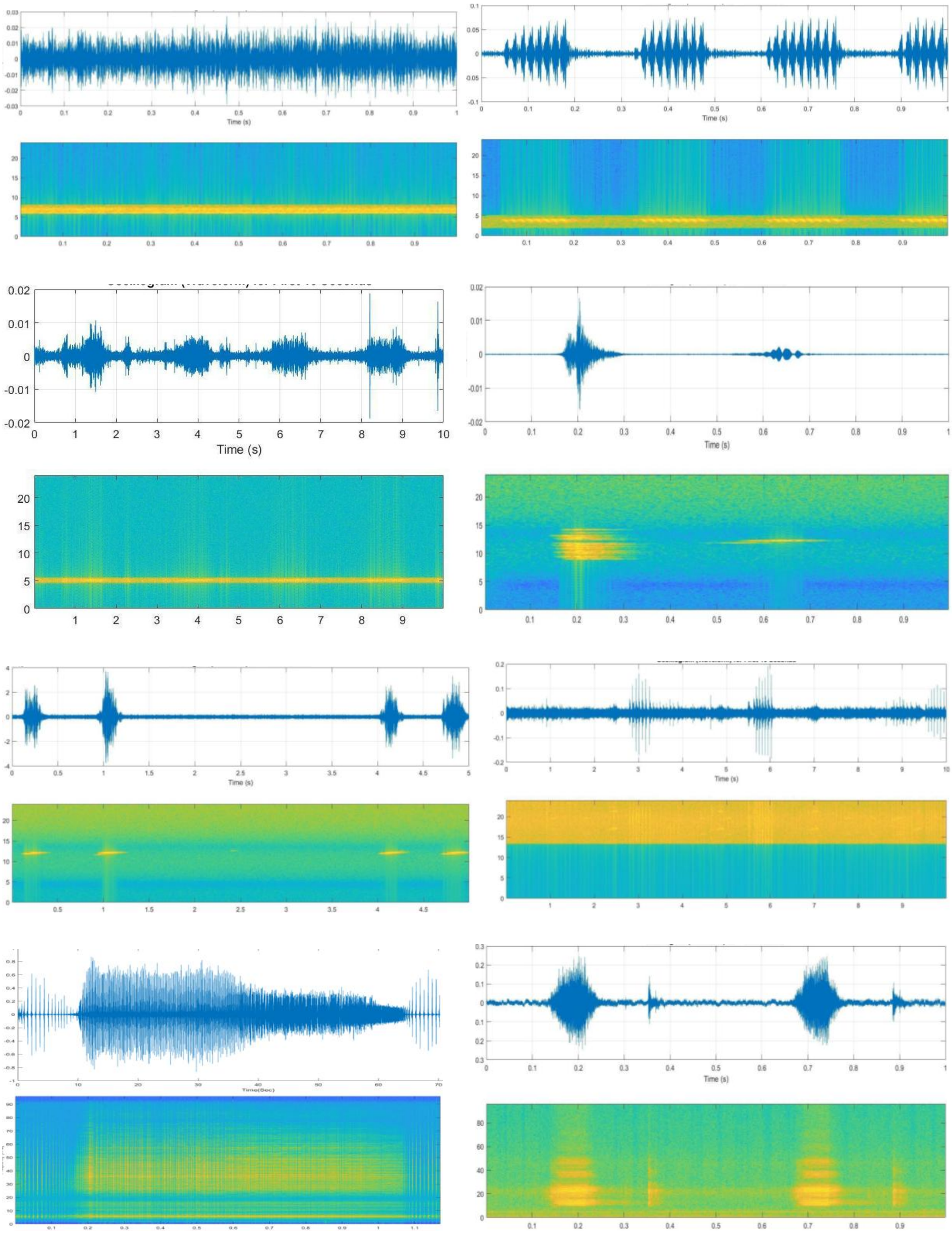

**Figure S1.** Oscillogram and spectrogram of each call type from ID1-ID8. Count clock wise (left -right- left).

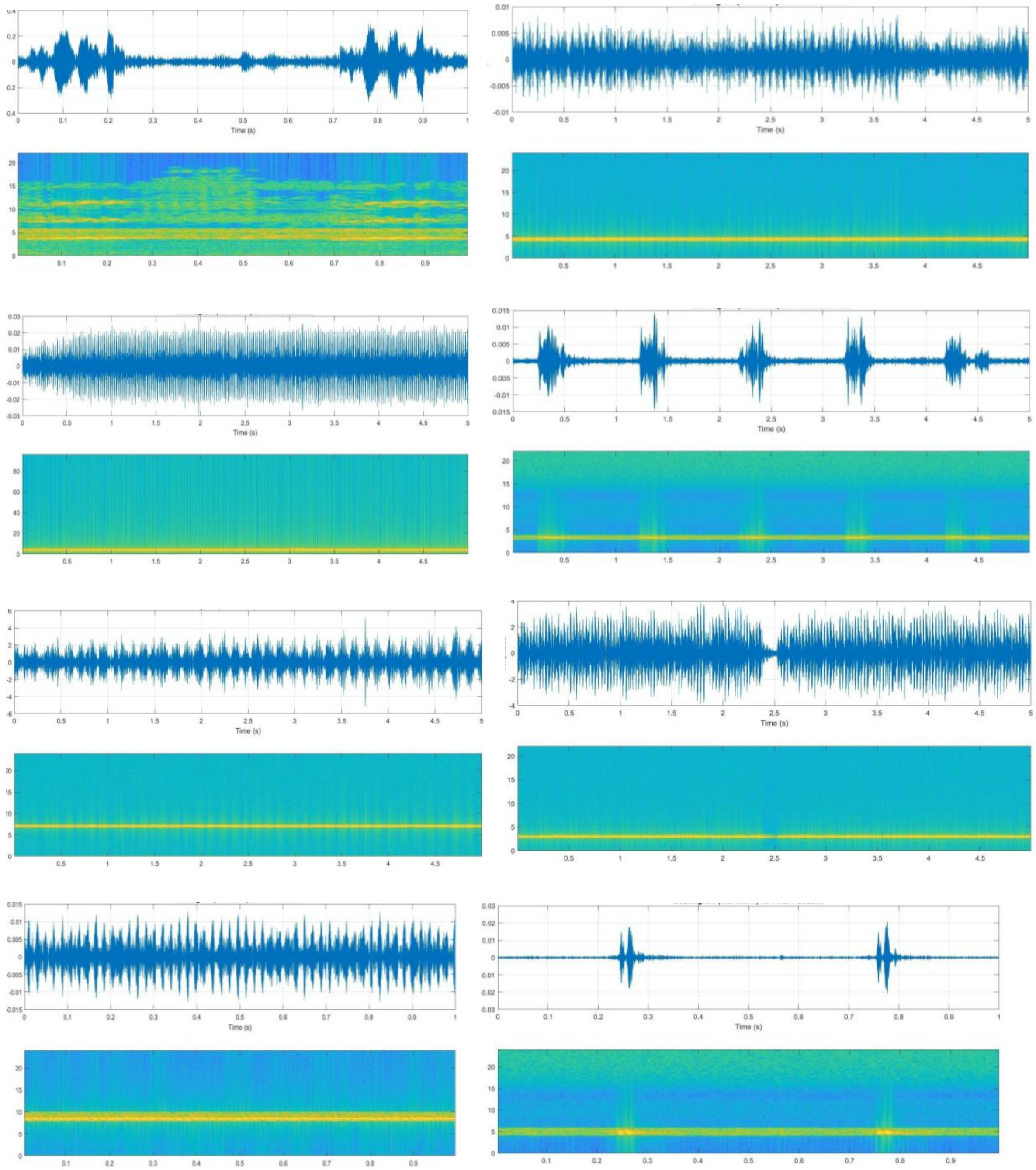

**Figure S2.** Oscillogram and spectrogram of each call type from ID9-ID16.

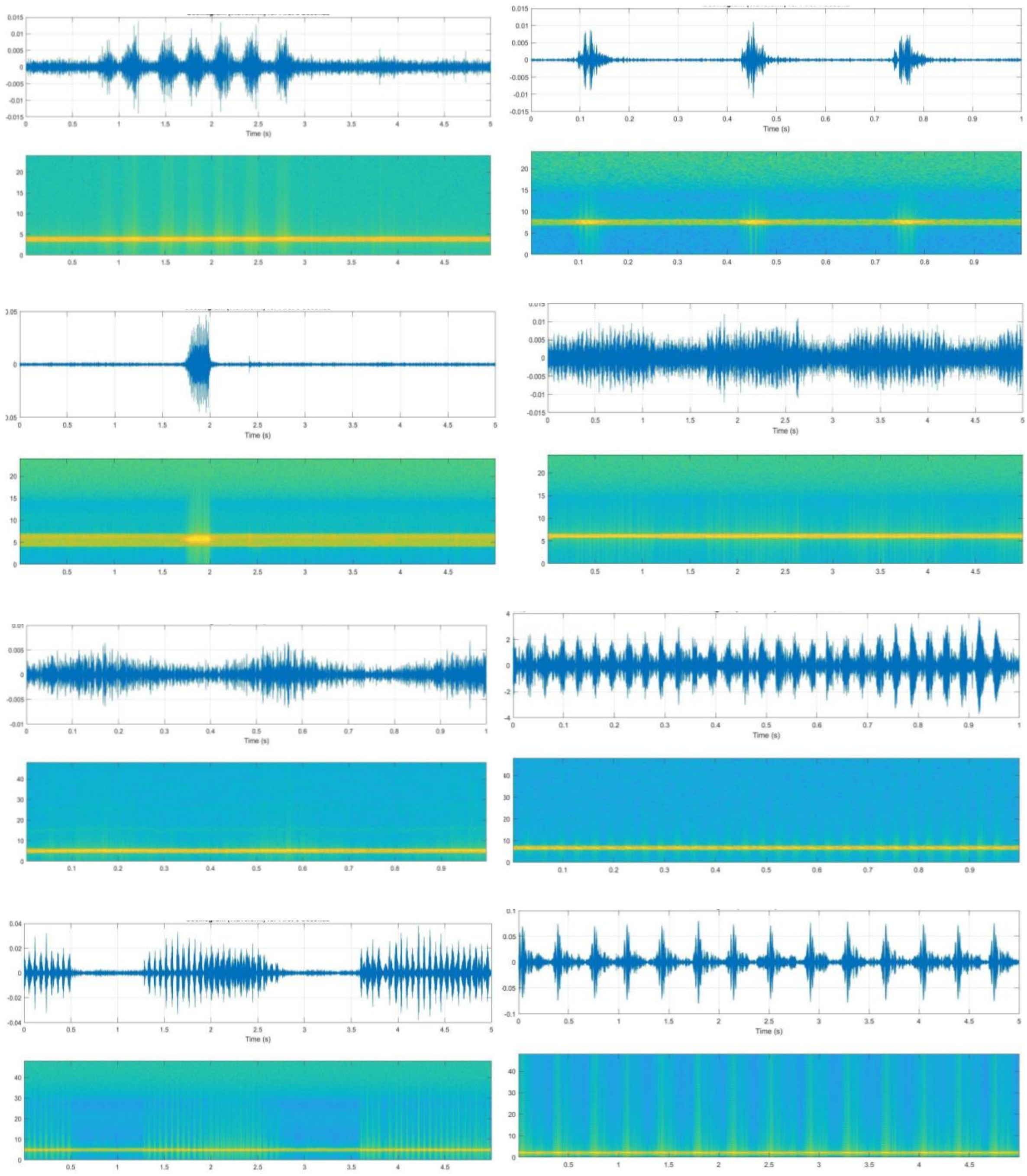


**Figure S3.** Oscillogram and spectrogram of each call type from ID17-ID24.

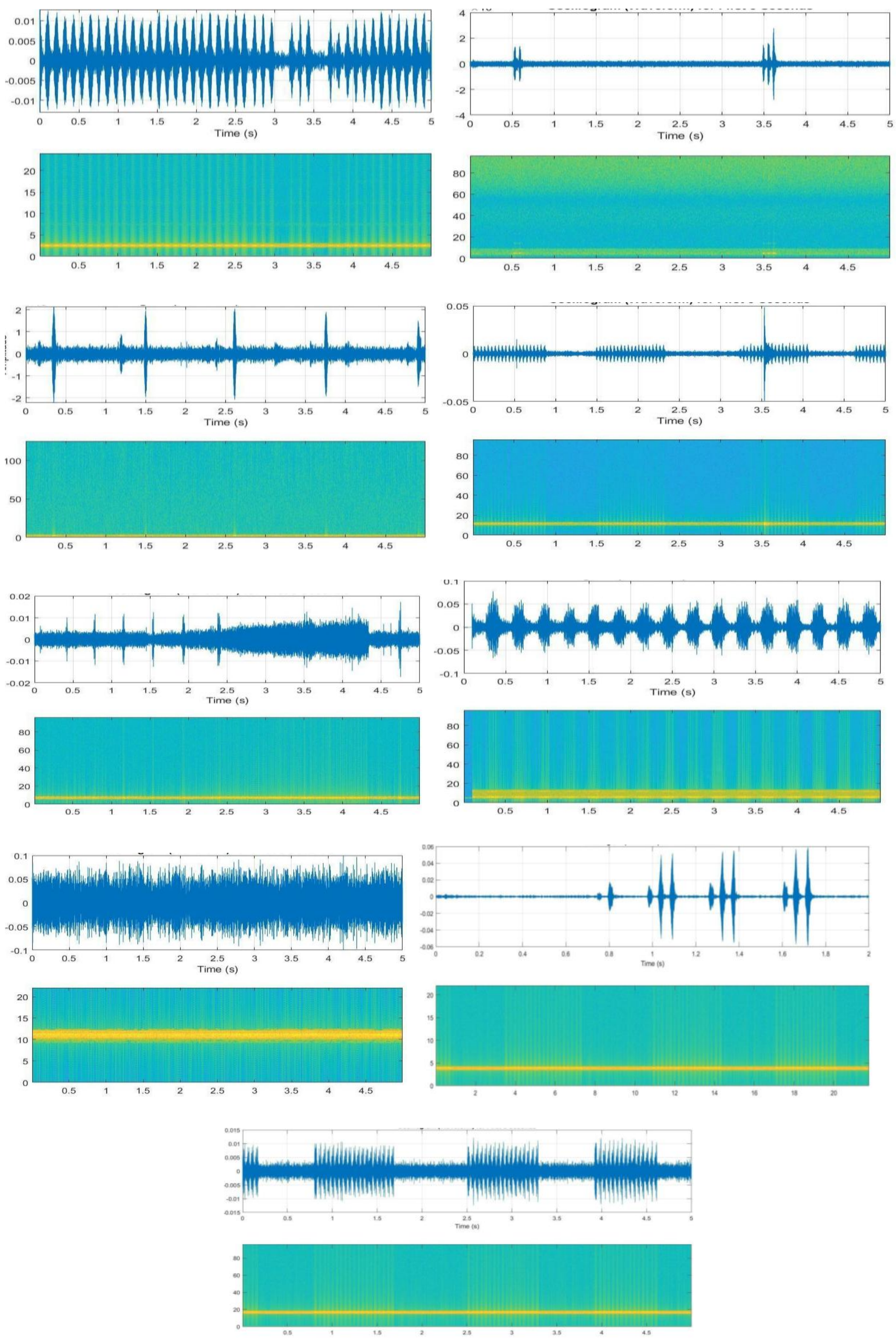


**Figure S4.** Oscillogram and spectrogram of each call type from ID 25-ID33

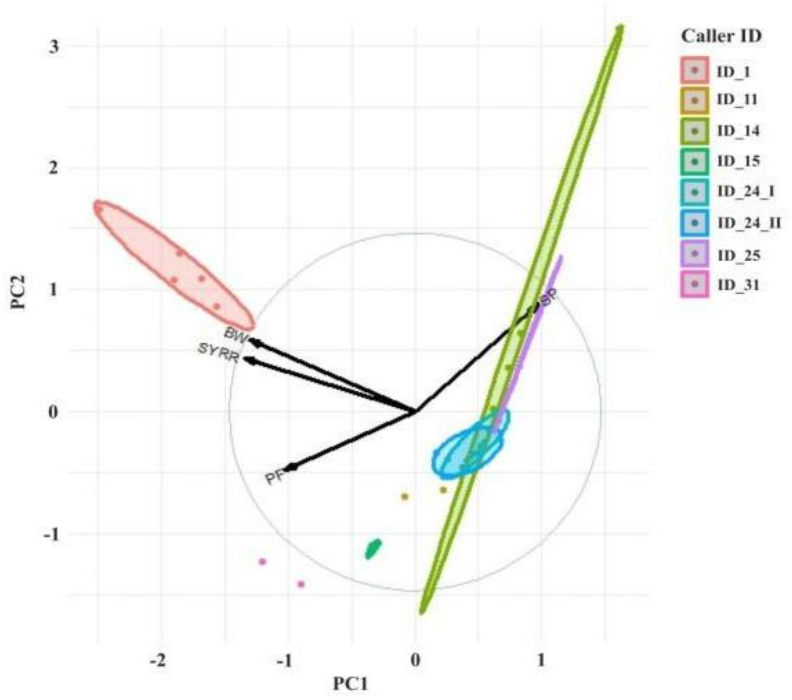


**Figure S5.** PCA biplot showing the direction of the axes representing different variables of all triller call types.

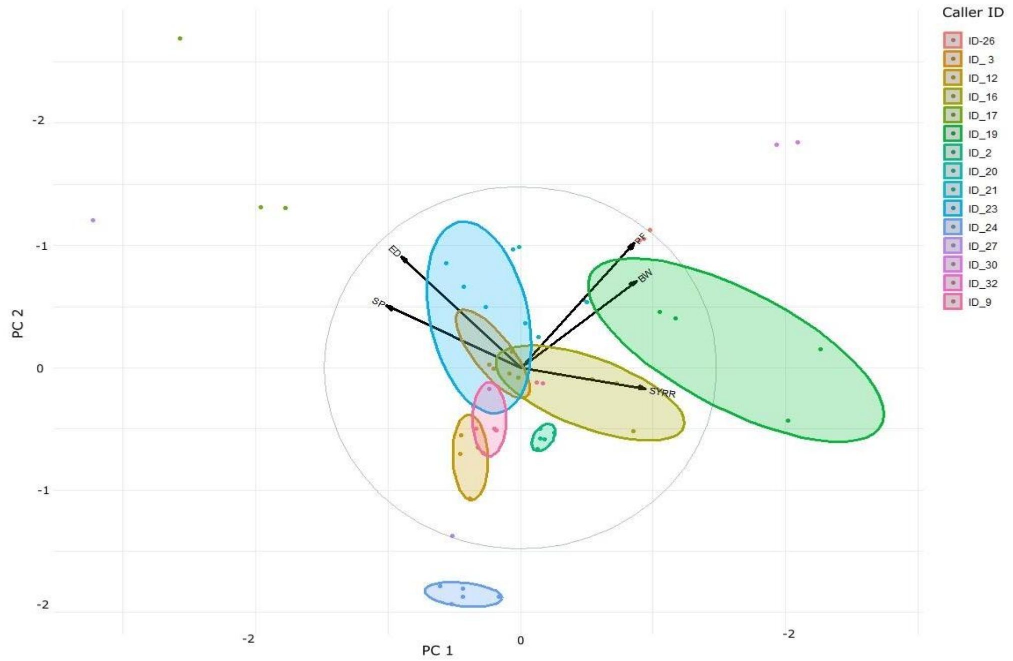


**Figure S6.** PCA biplot showing the direction of the axes representing different variables of all 3 high frequency chirper call types

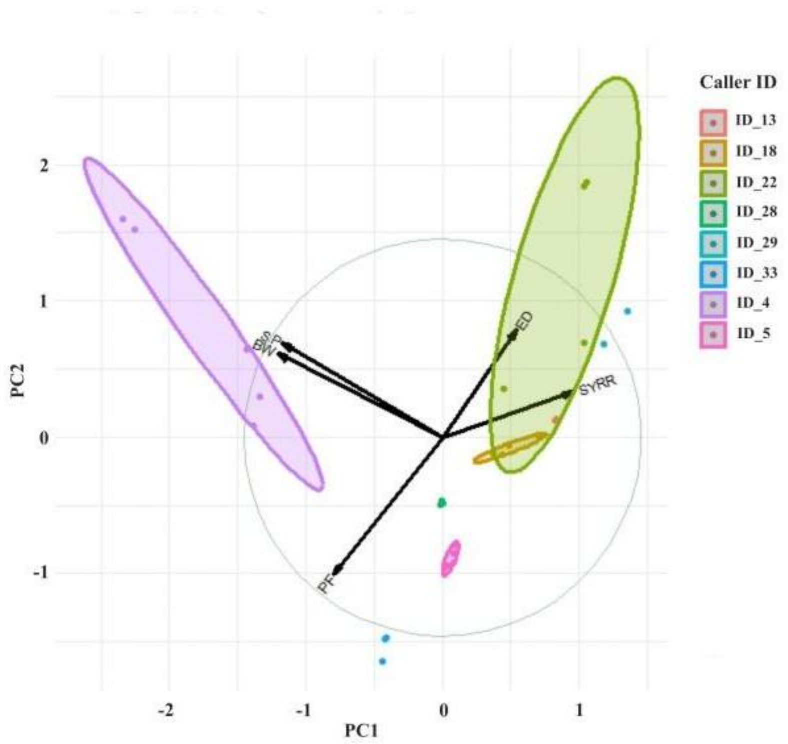


**Figure S7.** PCA biplot showing the direction of the axes representing different variables of all 7 low frequency chirper call types.
